# SOX9 is part of a combinatorial marker that reveals early development and embryological origins of the mouse brown adipose tissue depots

**DOI:** 10.1101/2025.07.16.665135

**Authors:** Ava E. Brent, Rika Chan, Vaishnavi Sirkay, Brian R. Morton, Jennifer H. Mansfield

## Abstract

Brown adipose tissue (BAT) is a mammalian thermogenic tissue that helps to maintain body temperature and provides important metabolic functions. Most brown adipocytes are located in structures called depots, and the development of depots is incompletely understood. Here, we define the embryonic positions in which the three major mouse BAT depots arise using a combination of morphological landmarks and expression of SOX9, a transcription factor that we find, by immunofluorescence and genetic lineage tracing, marks brown adipocyte progenitors from around the time of their exit from dermomyotome. Lineage tracing is also used to confirm the origin of the largest depot, interscapular BAT, in *Pax7-*positive dermomyotome of somites, and to show that the scapular BAT has a similar embryological origin, while cervical BAT arises from a somitic source outside of the *Pax7* lineage. Finally, using *Sox9* as part of a novel combinatorial marker, we profile the transcriptome of early brown adipocyte progenitors and pre-adipocytes and model their trajectory, revealing pathways including the paracrine signaling environment, in which brown adipocytes of the major depots likely arise. Our results apply to understanding the earliest stages of development of this novel mammalian tissue.

## Introduction

Brown adipose tissue (BAT) is a thermogenic tissue unique to mammals. BAT is the primary contributor to non-shivering thermogenesis in most species; it maintains body temperature, including during cold stress, and raises body temperature during the febrile response and arousal from hibernation (reviewed in (Cannon & Nedergaard, 2004; Oelkrug et al., 2015)). BAT is increasingly appreciated to have metabolic and endocrine roles as well. Active brown adipocytes oxidize lipids stored in intercellular droplets, and use the energy to generate heat by uncoupling the mitochondrial proton gradient from ATP synthesis. Thus, BAT activity also reduces overall metabolic efficiency, and its activity and expansion, which are centrally controlled, are thought to impact many aspects of physiology and medical conditions including diabetes and obesity. In adult humans, the BAT depots, although far smaller than those of infants, are estimated to account for about 5% of the basal metabolic rate (van Marken Lichtenbelt & Schrauwen, 2011).

Brown adipocytes are distributed within white adipose tissue throughout the body, and in small deposits around some visceral organs and blood vessels. However, most brown adipocytes are located in large, bilateral depots in the anterior trunk and neck, in proximity to the heart and major vessels. The BAT depots are believed to function in systemic warming. In both rodents and humans, interscapular BAT (iBAT) is the largest depot, located dorso-medial to the scapular blades and superficial to back and pectoral muscles. A large vessel (Sulzer’s vein in rodents) runs through iBAT and returns to the heart, carrying warmed blood for distribution to the body (Oelkrug et al., 2015). Scapular BAT (sBAT, also called axillary or subscapular BAT) forms deep to the scapular blades and is thought to warm blood returning to the heart via the subclavian and jugular veins. Cervical BAT (cBAT), located dorso-lateral to the cervical vertebrae, is near the common carotid and vertebral arteries that carry blood to the brain (Hachemi & U-Din, 2023; Sacks & Symonds, 2013).

The embryological origin and mechanisms through which BAT arises are not fully understood. Genetic lineage tracing in mice showed that the iBAT depot is derived from the dermomyotome compartment of somites. Somites are transient embryonic structures that generate musculoskeletal derivatives including most skeletal muscle, as well as the axial skeleton and its associated connective tissues (reviewed in Brent & Tabin, 2002). Somites form as bilateral epithelial balls flanking the midline, and subsequently divide into sclerotome, a ventral compartment that gives rise to cartilage, bone and tendon, and dermomyotome, a dorsal compartment that further segregates to produce the myotome, containing progenitors of skeletal muscles, and the dermatome, which contains dorsal dermis progenitors. Lineage tracing has shown that dermomyotome also gives rise to brown adipocytes (Lepper & Fan, 2010; Sanchez-Gurmaches & Guertin, 2014; Sebo et al., 2018; Sebo & Rodeheffer, 2019). It is likely that somites specifically at the axial levels where BAT depots arise (the anterior trunk and neck) produce most brown adipocytes of the major depots, although this has not been directly tested.

Perinatal and adult brown adipocytes of the iBAT have a history of expressing *Meox1,* a somite-specific marker, as well as *Pax7, Pax3, En1* and *Myf5,* which together indicate an origin in the central and dorsal dermomyotome (Atit et al., 2006a; Lepper & Fan, 2010; Seale et al., 2008; Sebo et al., 2018). This region also gives rise to epaxial skeletal muscle and dermis. sBAT has an expression history of *Pax3* and *Myf5* but not *En1,* suggesting a dermomyotome origin outside of the *En1*-expressing epaxial region (Ahmed et al., 2017; Atit et al., 2006a; Sanchez-Gurmaches & Guertin, 2014). Most cBAT adipocytes are labelled by *Pax3-Cre*, but unlike iBAT or sBAT, only about 60% are labelled with *Myf5-Cre* (Sanchez-Gurmaches & Guertin, 2014). Thus, it is accepted that brown adipocytes of the major BAT depots largely or entirely arise from dermomyotome although other sources are possible, including sources outside of somites.

The timing and mechanisms of brown adipocyte lineage commitment remain incompletely understood, especially during their embryonic origin *in vivo*. This is due in part to a lack of early molecular markers, although much progress has been made. The lineage labeling described above suggests that brown adipocytes, skeletal muscle and dorsal dermis share a common progenitor in dermomyotome. Of these three cell types, dermal fibroblasts make the earliest lineage commitment, based on the timing with which they extinguish *Pax7* expression (Lepper & Fan, 2010) and supported by scRNA-Seq analysis of the *Pax7* lineage (Fung et al., 2022). Dermal fibroblast fate is promoted by Wnt signaling from the overlying epidermis (Atit et al., 2006a). Brown adipocyte progenitors of iBAT extinguish *Pax7* later than dermis, but prior to embryonic day (E)12.5 (Lepper & Fan, 2010). GATA6 was recently found to promote brown adipocyte fate both during embryonic depot formation and during differentiation of iPS cells (Jun et al., 2023; Rao et al., 2023). Reciprocally antagonistic transcriptional circuits are involved in the specification of brown adipocyte fate vs. skeletal muscle fates (reviewed in (W. Wang & Seale, 2016)).

Brown adipocyte progenitors of the mouse iBAT depot have been detected *in vivo* as early as E12.5 by whole-mount mRNA *in situ* hybridization for adipocyte markers *Pparg*, *Cepba*, *Fabp4* and section immunofluorescence for GATA6 (Jun et al., 2023; Mayeuf-Louchart et al., 2019; Rao et al., 2023). scRNA-seq showed that *Gata6*-positive brown adipocytes derive from a multipotent fibroblast progenitor population present at E11.5-E12.5 (Jun et al., 2023; Rao et al., 2023). This work significantly advanced understanding of early iBAT development. However, sBAT and cBAT are less studied, and the earliest stages of BAT depot formation have not been visualized for any of the major depots. Characterizing the spatial and temporal emergence of brown adipocyte progenitors is necessary to understand the environments in which the depots develop, and the local cell and tissue interactions that regulate their origin *in vivo*.

Here, we define the positions in which the three major BAT depots arise relative to earlier-forming musculoskeletal landmarks. We show that all three are somite-derived based on their expression history of *Meox1*. Further, we show that the transcription factor SOX9 marks brown adipocyte progenitors and brown pre-adipocytes in all three depots, with an expression pattern that initiates before E11.5 in iBAT and sBAT and earlier in cBAT. This novel early brown adipocyte progenitor marker, visualized both by immunofluorescence and with conditional genetic lineage tracing, reveals that *Sox9* expression is activated in brown adipocyte progenitors earlier than other known markers, and thus facilitates visualization of early stages of depot development. *Pax7* conditional lineage labeling, in combination with SOX9 expression, can be used to visualize the positions of early brown adipocyte precursors in the iBAT and sBAT forming regions. In contrast, the cBAT arises outside of the *Pax7* dermomyotome domain and activates *Sox9* earlier, suggesting a distinct developmental origin. scRNA-Seq and trajectory analysis indicate that brown adipocyte progenitors are transducing signals relevant to the local environment and identify candidate drivers of brown adipocyte formation *in vivo*.

## Materials and Methods

### Mouse Lines, Breeding and Tamoxifen delivery

The following mouse alleles were used: *Rosa26*^tdTomat*o*^ Cre reporter*: B6.Cg-Gt(ROSA)26Sortm9(CAG-tdTomato)Hze/* (Madisen et al., 2010), Jax strain #007909; *Meox1*^Cre^ *: Meox1*^tm 1(cre)Jpa^ (Jukkola et al., 2005)*; Pax ^Cre/ERT2^*: B6.Cg-*Pax7^tm1(cre/ERT2)Gaka^*/J (Murphy et al., 2011), Jax strain # 017763; *Sox9* ^Cre/ERT2^: C57BL/6-*Sox9^em1(cre/ERT2)Tchn^*/J (Xu et al., 2015), Jax strain #035092 . To generate embryos, males bearing a *Cre* allele were crossed to females homozygous for the reporter transgene. Embryonic age was assigned as 0.5 day on the morning of detection of a vaginal plug. For conditional lineage labeling, a single intraperitoneal injection of 2mg tamoxifen (Sigma T5648, dissolved in 100µL sterile corn oil) was delivered to gestating females. All procedures were performed in accordance with the NIH Guide for Care and Use of Laboratory Animals and approved by the Columbia University IACUC.

### Genotyping

Genotyping was performed on tail biopsies or embryonic yolk sacs using GoTaq PCR mix (Promega) following manufacturer’s instructions. Primer sequences used are (from 5ʹ to 3ʹ): Cre: forward GCGGTCTGGCAGTAAAAACTATC, reverse GTGAA ACAGCATTGCTGTCACTT; tdTOMATO/RFP: forward CTG TTCCTGTACGGCATGG, reverse GGCATTAAAGCAGCGTA TCC.

### Immunofluorescence

Immunostaining was performed following previously described protocols (Holzman et al., 2018; McGlinn et al., 2019). Briefly, embryos were fixed overnight in 4% paraformaldehyde at 4°C and embedded in OCT. Sections of 12-15μm were cut, dried at 37°C, and frozen at −80°C. After air drying and washing in PBS, sections were permeabilized and blocked in 0.3% Triton-X/5% normal serum/PBS for 1 h. Primary antibodies were diluted in blocking solution and incubated overnight at 4°C. Antibody catalog numbers and dilutions are given in **Table S1.**

After washing in PBS, slides were incubated 3 h at room temperature in whole IgG secondary antibodies conjugated to Cy5, Alexa Fluor 488, Alexa Fluor 594 or Alexa Fluor 647 (Jackson Immunoresearch) at 1:200 or 1:400. Slides were stained with DAPI and mounted in Prolong Diamond (Invitrogen). Confocal images were captured on a Nikon A1 confocal unit attached to a Nikon Eclipse Ti2 microscope.

### Hematoxylin and Eosin Staining

Staining was performed on OCT cryosections or paraffin sections with an H & E kit (Abcam ab245880) following manufacturer’s instructions. Slides were mounted in gelvatol (McGlinn et al., 2019).

### mRNA *in situ* hybridization

mRNA in situ hybridization with a *Cepba* probe was performed as previously described (Chen et al., 2013). The *Cebpa* probe was generated by PCR-amplification from mouse trunk cDNA using the following primers: Fwd: 5’-GGAGTTGACCAGTGACAATG-3’ Rev: 5’CATTCTCCATGAACTCACCC-3’, subcloned into the p-Drive vector (Qiagen 231124) and verified by Sanger sequencing. The cDNA insert was PCR amplified and used as a template for antisense RNA probe transcription using a digoxigenin RNA labeling kit (Roche 11175025910).

### scRNA-Seq: Cell preparation

Embryos for scRNA-Seq were harvested at E11.5, E12.5 and E13.5. Trunks containing the segments C3-T2 (third cervical-second thoracic) were dissected in sterile PBS to remove surrounding tissues including the thoracic organs and forelimbs. In addition to somites and their derivatives, collected samples included the ventral body wall, epidermis and neural tissue from the C3-T2 axial level. Cells were dissociated by incubation in TrypLE Express (Gibco 12605010) for 15 minutes at 37°C, gently pipetting with a siliconized wide bore pipette every 5 minutes. All steps after dissociation were performed on ice. Dissociated cells were filtered through a FlowMi Cell Strainer (Sigma BAH136800040), and centrifuged at 184 RCF for 2 minutes at 4°C. Cells were resuspended in DMEM/20% FBS, and preparations from 3 littermates were then pooled at each timepoint. Cell concentration was estimated with a hemocytometer, and cells were centrifuged again as above and resuspended in DMEM/20% FBS at approximately 10^6^ cells/mL.

### scRNA-Seq: Sequencing and analysis

Sequencing and analysis were performed by the Columbia University Sulzberger Genome Center Single Cell Analysis Core facility. Cell viability and concentration was confirmed by the facility using Trypan blue and a Countess II cell counter. Samples were between 84-91% viable. Cells were encapsulated and libraries produced using the using 10x Genomics Chromium method (Zheng et al., 2017) with a Chromium Single Cell 3’ Reagent kit (version 3.1).

Sequencing was performed on an Illumina NovaSeqX Plus. FastQ files were processed and aligned to the mouse reference transcriptome (GRCm39-2024-A) using Cell Ranger 8.0.1 software. Numbers of cells in each timepoint sample (each of which contained cells from 3 littermates) were as follows: E13.5, 12,732 cells; E12.5 8,451 cells; E11.5 7,647 cells.

Analysis was performed using Anndata (Virshup et al., 2021)) and Scanpy (Wolf et al., 2018)). Sample files were concatenated and quality control filters were applied: cells with <200 expressed genes were removed (129 cells total, leaving the following numbers of cells in each sample: E13.5, 12,659 cells (73 cells removed); E12.5 8,418 cells (33 cells removed); E11.5 7,624 cells (23 cells removed) and filtering out genes expressed in <3 cells (10,636 genes). Cells with mitochondrial gene expression exceeding a threshold % were also removed (for E13.5, 17%, E12.5, 11%, E11.5, 10%). Scrublet was used to remove doublets (parameters: min counts = 2; min cells = 3; min gene variability = 85%; threshold set to the default (Wolock et al., 2019) (E13.5, 122 doublets removed; E12.5, 98 doublets removed; E11.5, 164 doublets removed).

Data were then normalized and logarithmized, the regress_out function was applied on total counts per cell, and data were scaled to unit variance. Principal components analysis (PCA) was performed, followed by dimension reduction with UMAP (McInnes et al., 2018). The Leiden method was used to generate cell clusters (Blondel et al., 2008). The bioinformatic work above was performed by the Single Cell Analysis Core; code is available on request. Differential expression analysis was performed using raw counts in Scanpy or in Loupe Browser 8.0 following export of Scanpy files to loupe format (code available on request), in order to identify marker genes for each sample. Leiden resolution was selected based on analysis of known marker genes and expected cell types. scRNA-seq data will be released at NCBI GEO upon acceptance for publication.

For RNA velocity analysis, spliced and unspliced transcripts were generated using Velocyto by the Sulzberger Genome Center(La Manno et al., 2018). Spliced, unspliced, and ambiguous matrices were generated and integrated as different layers into the original Anndata file. The Anndata file was subsetted to include only *Pdgfra+* clusters (see text), and the resulting file had 10879 cells and 18973 genes. scVelo was used to analyze RNA velocity data and the dynamical mode was used to estimate velocities (Bergen et al., 2020). Cellrank2 was used to estimate differentiation trajectories, infer cells’ fate probabilities, and visualize gene expression trends based on two kernels: experimental timepoints (RealTimeKernel) and RNA velocity (VelocityKernel)(Weiler et al., 2024). The two kernels were combined and given an equal weight of 0.5.

Pathways analysis was performed by searching for statistically over-represented GO Biological Process terms with FDR<.05 in Panther v19.0 (Mi et al., 2019; Thomas et al., 2022).

## Results

The anatomical positions of the mature mouse iBAT, sBAT, and cBAT depots along the AP and DV axes are schematized in **Figure S1A**. The depots can be visualized in tissue sections by immunofluorescence for PPARγ by E14.5 (**Figure S1B**), and the first signs of differentiation, such as lipid droplet formation, are detectable by E15.5 (Mayeuf-Louchart et al., 2019). mRNAs for committed pre-adipocyte markers have been detected earlier than proteins, at E12.5 by whole-mount *in situ* hybridization on iBAT (Mayeuf-Louchart et al., 2019), but not in section.

In order to characterize the anatomical positions of depots from their earliest appearance, and relative to earlier-forming musculoskeletal structures, we used section mRNA *in situ* hybridization for *Cepba* **(Figure S1D-F)**. All three depots could be visualized by strong *Cebpa* mRNA signal at E15.5, and faint signal at E13.5. At both stages, the sBAT depot appeared as a triangular region surrounded on all sides by muscle: it was bordered dorsally by the rhomboid, laterally by the levator scapula and medially by the epaxial muscle columns. A ventral extension of the sBAT was separated from the main sBAT triangle by the levator scapula and bounded laterally by the subscapularis muscle (**Figure S1D**). Collectively, this places the main sBAT triangle between the epaxial muscle columns and the hypaxial myotome-derived deep pectoral girdle muscles, and its ventral extension between deep and superficial pectoral girdle muscles (**Figure S1D,F**; for pectoral girdle muscle identification and classification, see (Valasek et al., 2010, 2011)). The cBAT depot was located lateral to the neural arch, fully enclosed by epaxial muscles derived from the medial and intermediate epaxial muscle columns (**Figure S1D,F**; for epaxial column formation and derivatives, see (Deries et al., 2010; Mekonen et al., 2016)). The iBAT depot was located superficial to the rhomboid and trapezius muscles and medial to the scapula (**Figure S1E-F**). The consistency of BAT depot location relative to surrounding structures over developmental time suggests that they develop largely in place, like the epaxial muscles. Thus, positions of depot emergence may be predicted by visualizing the earlier-developing epaxial and pectoral girdle muscles. Interestingly, at E12.5, a cluster of mesenchyme was present in the future position of each depot, relative to the same musculoskeletal landmarks, but was negative for *Cebpa* mRNA (**Figure S1D-E**). This mesenchyme likely includes brown adipocyte precursors, as verified for the iBAT depot (Jun et al., 2023; Rao et al., 2023), but not for the other depots.

We next sought to identify a marker that would label brown adipocyte precursors from their earliest emergence. We observed expression of the transcription factor SOX9 in BAT depots and thus proceeded to characterize it further. In somites, the role of SOX9 is best described in the sclerotome compartment, where it is essential for the specification and differentiation of sclerotome cells into the cartilage of the vertebrae and ribs (Lefebvre et al., 2019). SOX9 is additionally expressed transiently in sclerotome-derived axial tendon progenitors (Asou et al., 2002; Sugimoto et al., 2012), and in the muscle connective tissue (MCT) of epaxial and hypaxial muscles (pink and blue arrows in **Figure 1Eii**).

**Figure 1.**
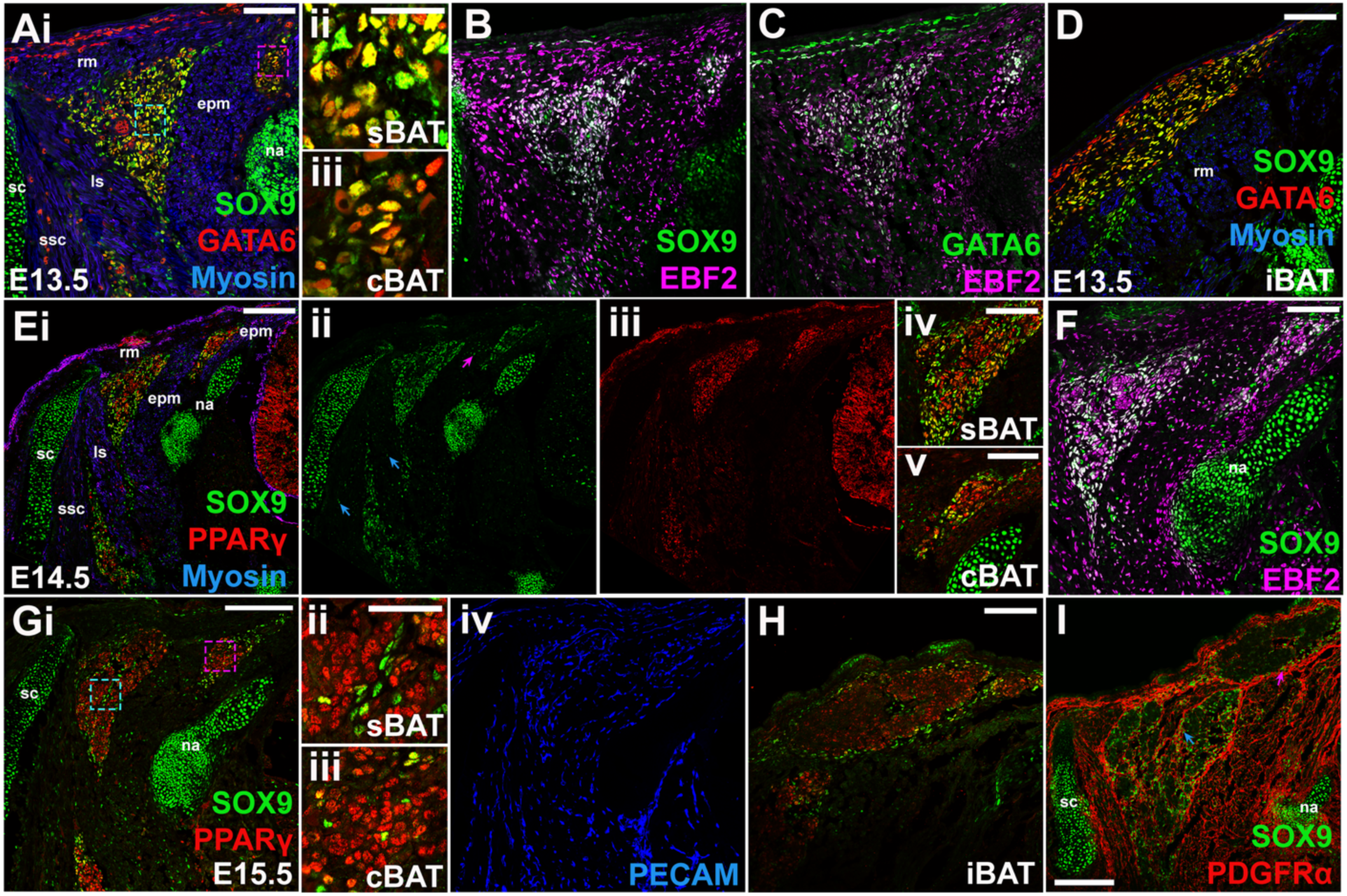
SOX9 marks brown adipocyte progenitors and brown pre-adipocytes. Co-expression of SOX9 and GATA6 (A, D), SOX9 and EBF2 (B), and GATA6 and EBF2 (C), in alternate transverse sections through the cervical region of E13.5 embryos. sBAT and cBAT depots are visible in (A-C), iBAT in (D). Skeletal muscle myosin shown in blue in (A) and (D). (Aii) and (Aiii) show higher magnification of regions marked by, respectively, the blue box (sBAT) and pink box (cBAT) in (Ai). Co-expression of SOX9 and PPARγ (Ei), SOX9 alone (Eii), and PPARγ alone (Eiii) in transverse sections through the cervical region of E14.5 embryos. Skeletal muscle is marked by myosin expression in (Ei). Higher magnification view of sBAT and cBAT from (Ei), shown in (Eiv) and (Ev). Arrows in Eii indicate expression of Sox9 in muscle connective tissue of epaxial (pink) and hypaxial (blue) muscles. (F) Co-expression of SOX9 and EBF2 in sBAT and cBAT depots in a transverse section through the cervical region of E14.5 embryos. (Gi) Co-expression of SOX9 and PPARγ in a transverse section through the cervical region of an E15.5 embryo. (Gii) and (Giii) show higher magnification of regions marked by, respectively, the blue box (sBAT) and pink box (cBAT) in (Gi). (Giv) PECAM staining in same section shown in (Gi) reveals blood vessels. (H) Co-expression of SOX9 and PPARγ in a transverse section through the thoracic region of the same embryo shown in (G). (I) Co-expression of SOX9 and PDGFRα in a transverse section through the thoracic region in an alternate section to that shown in (H). Arrows indicate SOX9 and PDGFRα co-expression in fibroblasts of the iBAT (pink arrow) and sBAT (blue arrow). Scale bars: 200µm: (Ei), (G); 100µm: (Ai), (D), (Ei), (F), (H), (I); 25µm: (Aii), (Gii). Abbreviations: epm, epaxial muscles; ls, levator scapula; na, neural arch; rm, rhomboid major; sc, scapula; ssc, subscapularis muscle.

In transverse sections through the cervical region (C4-C5) of E13.5 mouse embryos, we observed specific and robust SOX9 expression in the anatomical positions of the sBAT and cBAT depots (**Figure 1A**; compare positions to **Figure S1D,F**)). Further, SOX9 expression co-localized with GATA6, a transcription factor expressed in brown adipocyte progenitors of iBAT starting at E12.5 ((Jun et al., 2023; Rao et al., 2023) (**Figure 1A**)), and with EBF2, a transcription factor selectively expressed in brown versus white adipocytes and progenitors (W. Wang et al., 2014) but also broadly expressed in muscle connective tissue (MCT), other mesenchyme and some cartilage **(Figure 1B, C** and see (Holzman et al., 2021)). Transverse sections through the anterior thoracic region (T1) showed that SOX9 was also expressed in the anatomical position of the iBAT depot, where it also co-localized with GATA6 (**Figure 1D**; compare to **Figure S1E-F**) and EBF2 (not shown). Finally, in all three depots, SOX9 was co-expressed with HOXA5, which we previously reported as an early but not specific marker for brown adipocyte progenitors (**Figure S1C** and not shown; (Holzman et al., 2021)). Thus SOX9, GATA6, EBF2 and HOXA5 are co-expressed by brown adipocyte progenitors in all three depots.

Having confirmed that SOX9 marks brown adipocyte progenitors of all depots at E13.5, we followed expression as they matured. PPARγ expression is first detected in pre-adipoctyes of the BAT depots at E14.5, and many of these cells highly co-express SOX9 (**Figure 1 Ei-v**). However, in contrast to E13.5 where the majority of cells in all depots expressed SOX9 (**Figure 1A-D**), two populations were apparent: SOX9+, PPARγ+ cells and SOX9-, PPARγ+ cells (**Figure 1E**). Both populations expressed EBF2 (**Figure 1F**). This is consistent with the known persistence of PPARγ and EBF2 in pre-adipocytes and differentiating adipocytes, and suggests SOX9 becomes downregulated as cells begin to differentiate. Supporting this interpretation, the SOX9-, PPARγ+ cells appeared morphologically more advanced than the SOX9+ PPARγ+ cells. The former were organized into in clusters, resembling the structure of differentiated BAT, while the latter appeared as a loose mesenchyme like at E13.5 (**Figure 1Ei, F**). A similar distribution of SOX9+, PPARγ+ and SOX9-, PPARγ+ cells was seen in the E14.5 iBAT depot (not shown).

By E15.5, brown adipocytes have more fully adopted a clustered morphology and are surrounded by PDGFRα-expressing connective tissue (**Figure 1G-I**). In addition, the depots show high blood vessel infiltration, which is essential to BAT’s heat-producing function (**Figure 1Giv**). This tissue organization is maintained in the mature depots. At E15.5 nearly all brown adipocytes, marked by PPARγ, were SOX9- (**Figure 1Gi-iii; H)**. However, SOX9 expression persisted within PDGFRα+ fibroblasts of all three depots (arrows in **Figure 1I** and not shown); this population comprises connective tissue within and encasing the developing BAT depots. A similar program of expression was reported for GATA6, which becomes down-regulated in brown adipocyte progenitors as they transition to pre-adipocytes, but is retained in a fibroblast connective tissue population surrounding the iBAT (Jun et al., 2023; Rao et al., 2023).

PDGFRα+ SOX9+ fibroblasts are likely a source for new brown adipocytes. In adult BAT undergoing cold-induced expansion, PDGFRα+ fibroblasts serve as a source for new brown adipocytes (Y. H. Lee et al., 2015); embryonically, *Pdgfra+* progenitors give rise to most mesodermal cell types of mesenchymal origin, including brown adipoctyes (Jun et al., 2023; Rao et al., 2023 and references therein), but the specific contribution these cells within embryonic BAT connective tissue is not established.

Having identified SOX9 as a specific and early marker of brown adipocyte progenitors and pre-adipocytes, and as a persistent marker of BAT connective tissue, we next employed an inducible Cre lineage-labeling approach to define the timeline during which the brown adipocyte lineage activates *Sox9* expression. We used a tamoxifen-inducible *Sox9*^Cre-ERT2^ allele and *Rosa26*^tdTomat*o*^ Cre reporter (Madisen et al., 2010; Xu et al., 2015) to mark the descendants of *Sox9*-expressing cells. A single tamoxifen injection between E8.5 and E12.5 was used to induce recombination in the reporter transgene, and embryos were allowed to develop until E15.5, when PPARγ expression is robust in all three depots. We then performed immunofluorescence for RFP and PPARγ to identify the contribution of *Sox9*-descendant (RFP+) cells to each depot.

A prior report showed that Cre-ERT2 nuclear translocation is detectable in embryos 6 hours after maternal IP tamoxifen injection, robust by 24 hours, and undetectable again by 48 hours (Hayashi & McMahon, 2002). Thus, RFP should mark cells that, at the earliest, expressed *Sox9* about 6 hours after the injection, and throughout the window during which ERT-Cre remained nuclear. Consistent with this expectation, following control tamoxifen injections with this line during stages when *Sox9* is continuously expressed, RFP protein was detectable by 12 hours, and robust in SOX9+ cells by 18-24 hours post injection (**Fig S2A-G**). Based on all of this, we assumed 6-24 hours as a conservative range for the timeline with which, following tamoxifen injection, *Sox9*-expressing cells would be labelled (by nuclear Cre-ERT activity).

Tamoxifen was first delivered at E8.5 to label *Sox9*-expressing cells in early compartmentalized somites (as early as E8.75-E9.5 using the 6-24 hour estimate). We observed labeling of adipocytes in cBAT, but not in the sBAT or iBAT depots, revealing that a significant number of cBAT adipocyte progenitors expressed *Sox9* at this early stage (**Figure 2A**). Injection a day later, at E9.5, was expected to mark *Sox9* expressing cells as early as late-stage compartmentalized somites (as early as E9.75-E10.5). Virtually all of the cBAT adipocytes were labelled by this treatment, as were a few sBAT adipocytes, while the iBAT remained largely unlabeled (**Figure 2B**). Injection at E10.5, to mark cells that activate *Sox9* as early as E10.75-E11.5, a time when cells begin exiting the dermomyotome by EMT, labelled an increasing number of sBAT and iBAT adipocytes but a slightly decreased number of cBAT cells (**Figure 2C**). Finally, tamoxifen injection at E11.5 or E12.5 (expected cells that activate *Sox9* as early as E11.75-E12.5 and E12.75-E13.5, respectively) a yielded robust labeling of sBAT and iBAT. In contrast, labeling of cBAT adipocytes was reduced by injection at these later stages (**Figure 2D, E**). Together, this reveals that a substantial number of brown adipocytes in the iBAT and sBAT depot are expressing *Sox9* at or prior to E11.5, given that they are marked by E10.5 injection, and virtually all express *Sox9* by E12.5 given that they are marked by injection at E11.5. As described above, immunofluorescence confirmed that SOX9 remained expressed in brown adipocyte progenitors and pre-adipocytes in all BAT depots during E13.5-E14.5, until expression became restricted to BAT connective tissue fibroblasts at around E15.5 (**Figure 1G-I**).

**Figure 2.**
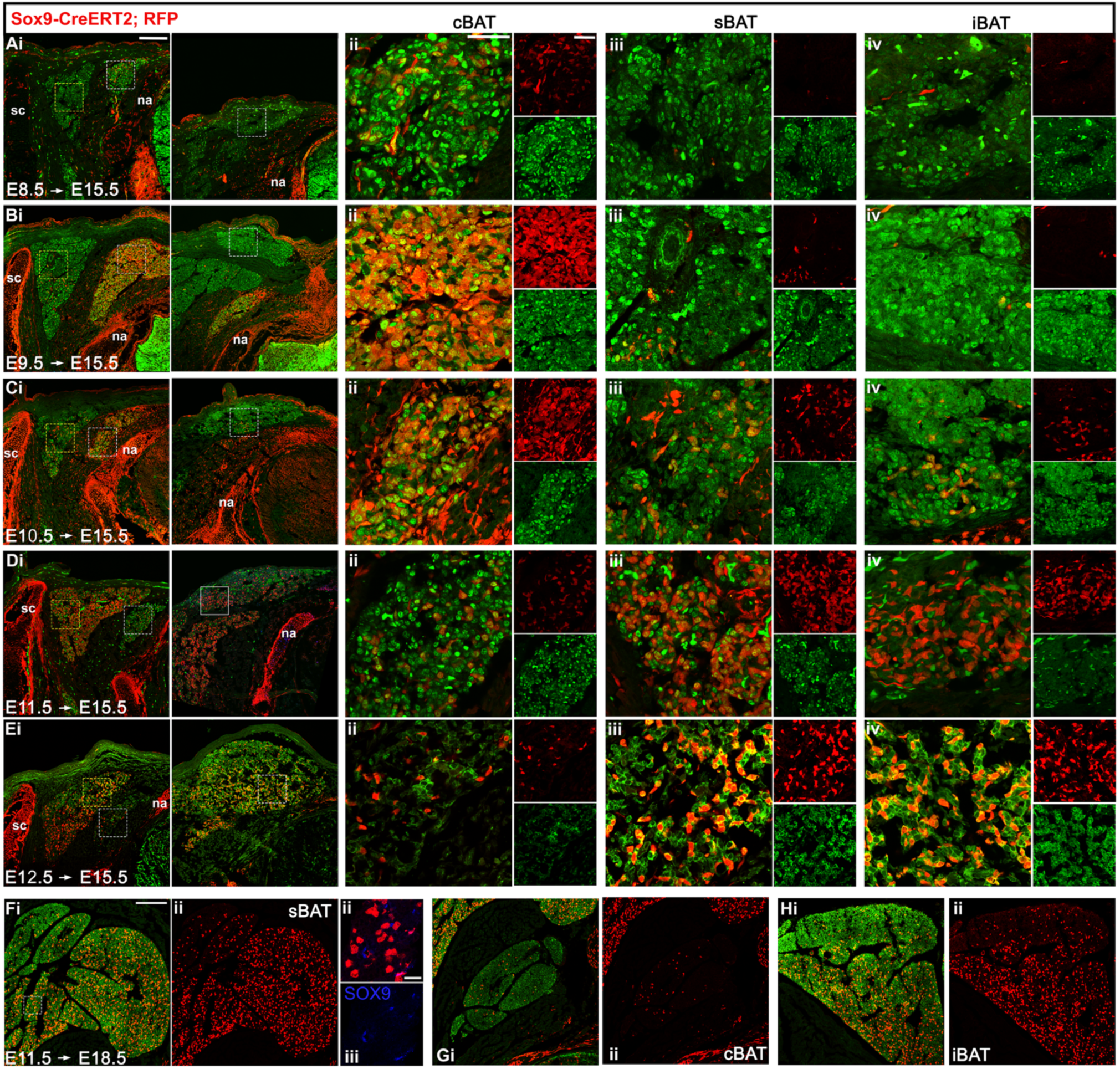
Conditional lineage labeling reveals depot-specific time-course for the activation of *Sox9* expression in brown adipocyte progenitors. Co-expression of PPARγ (for brown pre-adipocytes), and RFP (for the *Sox9* lineage) is shown in transverse view through either cervical sections (showing sBAT and cBAT, A-Ei, left panels) or thoracic sections (showing iBAT, A-Ei, right panels) from *Sox9^Cre/ERT2/+^; Rosa26^tdTomato/+^* embryos. Boxed regions in A-Ei are shown in closeup in A-Eii-iv, with depots as labelled; larger panels show an overlay and smaller panels show individual channels. The cBAT and iBAT closeup regions are indicated by white boxes in the left and right panels of A-Ei, respectively, and the sBAT closeup region is indicated a yellow box in Ai-Ei, left panel. A single injection of tamoxifen was given at the indicated stages, and embryos were harvested at E15.5. (F-H) Co-expression of PPARγ and RFP in transverse sections through each depot, as indicated, following tamoxifen injection at E11.5 and harvest at E18.5 to track the fate of *Sox9* lineage labelled cells following fetal BAT depot expansion. Scale bars: 200µm (Ai-Fi), 50µm (Aii-iv – Eii-iv), 25µm (Fii-iii). Abbreviations cBAT, cervical BAT; iBAT, interscapular BAT; na, neural arch; sBAT, scapular BAT; sc, scapula.

Together, these results defined a depot-specific range of stages during which *Sox9* expression is activated in brown adipocyte progenitors, schematized in **Figure S2M**. Notably, SOX9 expression in cBAT appeared more dynamic than in sBAT or iBAT. Combined SOX9 lineage and immunofluorescence analyses indicate that cBAT precursors express *Sox9* in compartmentalized somite stages (**Figure 2B, C**), and again in early cBAT depots from E13.5-E14.5 (**Figure1 A, E**). However, during the transition from compartmentalized somite to clearly demarcated cBAT depot, the number of cBAT *Sox9* expressing cells appears reduced (**Figure 2D, E**), suggesting the possibility of temporal waves of SOX9 expression unique to the cBAT.

While this time course defines the history of *Sox9* activation in adipocytes of mid-gestation depots, significant growth occurs between E15.5 and the end of gestation when depots become functional. Interestingly, tamoxifen injection at E11.5 labelled the majority of sBAT and iBAT adipocytes when observed either at E15.5 (**Figure 2D**) or E18.5 (**Figure 2F,H**). The relative consistency of labeling suggests that most adipocytes in mature depots are derived from cells expressing *Sox9+* as early as E11.75-E12.5. Finally, few cBAT adipocytes were labelled by injection at E11.5, observed at either E15.5 (**Figure 2D**) and E18.5 (**Figure 2G**).

By 15.5, when embryonic brown pre-adipocytes have commenced differentiation, SOX9 expression became restricted primarily to PDGFRα-positive BAT fibroblasts (**Figure 1G-I**). To determine the fate of SOX9+ fibroblasts, we injected tamoxifen at E15.5. As shown in **Figure S2I-L**, this produced substantial labeling of adipocytes in all three depots at E18.5. Further, like at E15.5, SOX9 expression at E18.5 remained mostly restricted to the PDGFRα-expressing connective tissue fibroblasts of BAT (**Figure S2J,L**). Together, this shows that SOX9+PDGFRα+ fibroblasts within BAT connective tissue are a source of new brown adipocytes during fetal BAT growth. This is consistent with other findings that BAT connective tissue fibroblasts can be a source of new adipocytes during fetal and post-natal growth (Jun et al., 2023; Y. H. Lee et al., 2015).

We additionally observed *Sox9*-descendant cells in other mesodermal lineages, which is relevant to the interpretations below. Following tamoxifen injection at E8.5, *Sox9* descendants made up a minor proportion of the sclerotome-derived cartilage of the vertebrae, and the scapula cartilage was only very sparsely labeled (**Figure 2A** and not shown). Injection of tamoxifen at E9.5 and later yielded labeling of *Sox9* descendants throughout all cartilage elements. SOX9 expressing muscle connective tissue was also well-labeled when tamoxifen was injected at E9.5-E11.5, but not later (**Figure 2B-E**). Importantly for our ability to distinguish BAT progenitors from other dermomyotome derivatives, *Sox9*^Cre-ERT2^ did not label skeletal muscle. A few dermal fibroblasts were labelled by early tamoxifen injections, but not by injection at E11.5 or later (**Figure 2A-E)**.

Together, the results above show that SOX9 expression marks brown adipocyte progenitors earlier than they have previously been observed *in vivo*. We therefore wished to use this novel marker to study early stages of depot formation. However, the relatively broad expression of SOX9 in cartilage and connective tissues as well as brown adipocyte progenitors still presented a challenge to interpretation. We hypothesized that SOX9 expression could be used in combination with other established markers, such as EBF2 and/or a lineage label for dermomyotome to distinguish brown adipocytes from other SOX9-expressing cell types. As described above, *Pax7*+ dermomyotome cells give rise to brown adipocytes. Thus, we next used a *Pax7^Cre-ERT2^* allele (Murphy et al., 2011) with the *Rosa26*^tdTomat*o*^ Cre reporter line to characterize the contribution of *Pax7*-expressing cells to each *Sox9+* lineage, and to establish a depot-specific time-course for when brown adipocyte progenitors express *Pax7*.

A single tamoxifen injection was performed between days E8.5 and E11.5, followed by IF for SOX9 and PPARγ at E14.5 (**Figure 3**). Overall, our results fully agreed with previous findings about the timing with which brown adipocytes of the iBAT extinguish *Pax7* (Lepper & Fan, 2010) and added depot-specific detail regarding the *Pax7* lineage. Tamoxifen injection at either E8.5 or E9.5 resulted in near-complete labeling of the sBAT and iBAT depots, indicating an origin for both in *Pax7-*expressing dermomyotome of compartmentalized somites (**Figure 3A-B**). Injection at E10.5 labelled a decreased number of adipocytes in both depots. In sBAT, unlabeled adipocytes were spatially restricted to the dorsolateral-most corner adjacent to the scapula (**Figure 3Ciii**, arrow), whereas in the iBAT, they were dispersed throughout the depot (**Figure 3Civ**). Injection at E11.5 did not label adipocytes in the iBAT, and only a very few, medial-most adipocytes in sBAT (**Figure 3D**). Taken together, our observations indicate that the sBAT and iBAT progenitors are specified from a *Pax7* expressing cell population labeled by injection between E8.5-E10.5 but not later, and suggest that sBAT brown adipocytes extinguish *Pax7*, and thus may exit dermomyotome, in a lateral to medial temporal order. In contrast to iBAT and sBAT, we never observed a contribution of *Pax7*-descendants to the cBAT depot (**Figure 3Aii-Dii**).

**Figure 3.**
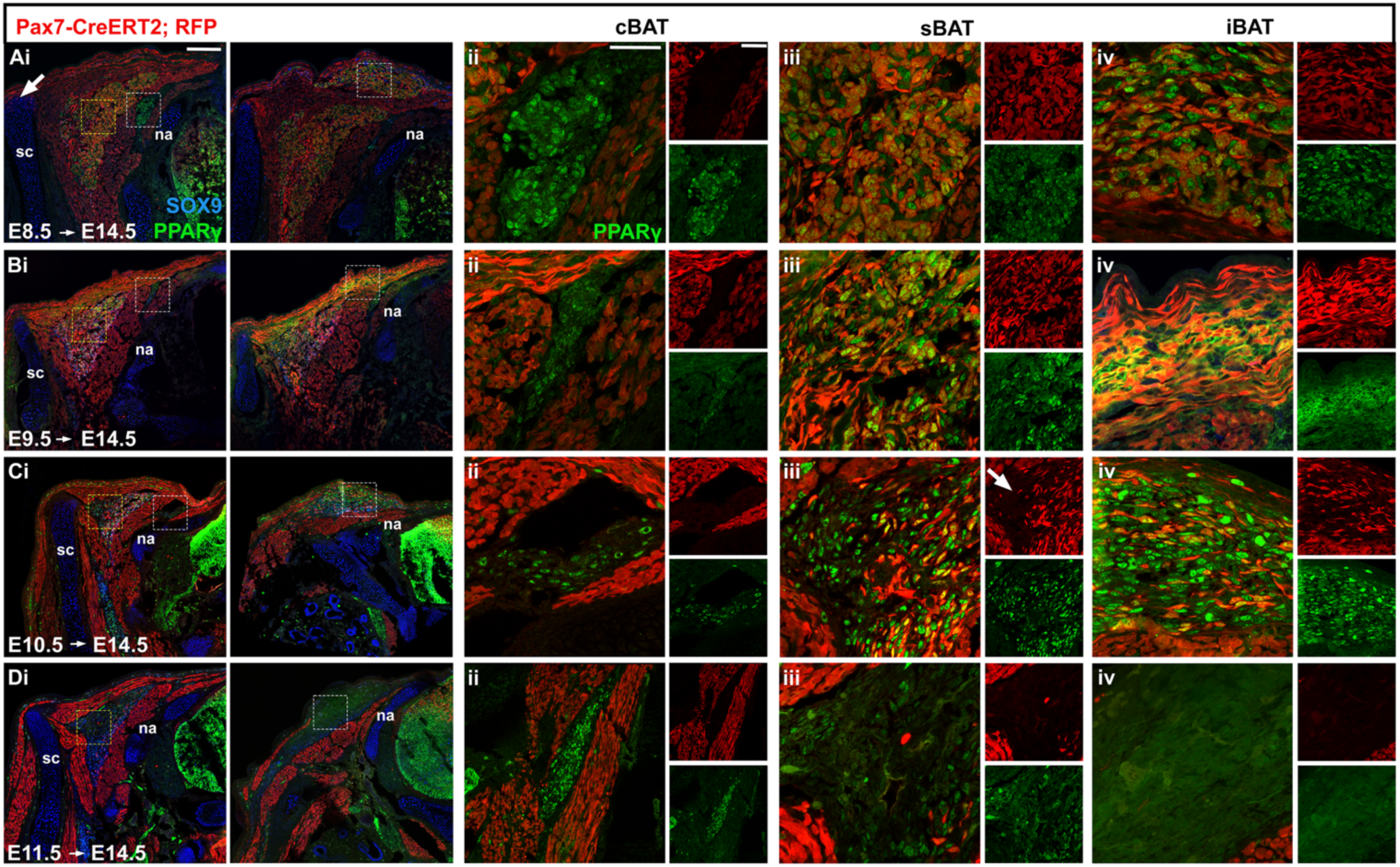
A depot-specific time course for *Pax7* expression shows a history of *Pax7* expression in iBAT and sBAT but not cBAT. Co-expression of PPARγ and SOX9 (for brown pre-adipocytes), and RFP (for the *Pax7* lineage) are shown in transverse view through either cervical sections (showing sBAT and cBAT) or thoracic sections (showing iBAT) regions in E14.5 *Pax7^Cre/ERT2/+^; Rosa26^tdTomato/+^* embryos. A single injection of tamoxifen was given at the indicated stages. (Ai-Di) show lower magnification views through cervical (left panel) or thoracic (right panel) segments of the same embryo. Higher magnification views shown in (A-D ii-iv) correspond to boxes in (Ai-Di) (yellow boxes indicate sBAT, white boxes indicate cBAT (left panels) and iBAT (right panels)). Note that the *Pax7* lineage label is absent from cBAT in A-D, but present as expected in skeletal muscle surrounding it. Scale bars: 200µm: (A-Di), 50µm: (A-Dii-iv). Abbreviations: cBAT, cervical BAT; iBAT, interscapular BAT; na, neural arch; sBAT, scapular BAT; sc, scapula.

The *Pax7* lineage label marked additional cell types, including skeletal muscle of the epaxial columns and pectoral girdle, and the dermis, as expected. We also observed a few cells labeled in the dorsal-medial border of the scapular blade after E8.5 but not later injections (**Figure 3Ai**, arrow) consistent with a dermomyotome (or *Pax3+*) origin for the medial scapular blade previously reported in chick and mouse, respectively (Huang et al., 2000; Valasek et al., 2010). Importantly, following injection at E9.5, the majority of SOX9-expressing, *Pax7*-descendant cells were found in the sBAT and iBAT, including both adipocytes and BAT connective tissue **(Figure S3Ai-iii** and data not shown). However, a small number of SOX9+, *Pax7* descendants were observed in a neighboring tissue, the MCT of the levator scapula and rhomboid muscles (both deep pectoral muscles, **Figure S3Ai, ii, iv**), but they were not observed in the MCT of the epaxial muscles or of the subscapularis (a superficial pectoral muscle) (**Figure S3Ai, ii, v**). The *Pax7* lineage did not contribute to vasculature of the BAT depots (**Figure S3B**) or to the tendons associated with the scapula or vertebrae (**Figure S3C**). Some tenascin-positive *Pax7* descendants were observed near the insertion sites of the rhomboid and levator scapulae on the medial-dorsal scapular blade, likely comprising myotendinous junctions of these deep pectoral girdle muscles **(Figure S3C**). Thus, the combination of *Pax7* lineage label induced at E9.5 with SOX9 expression allows us to distinguish iBAT and sBAT adipocyte and connective tissue progenitors from other SOX9 expressing cell types in the vicinity of BAT including most MCT, and all tendon or other connective tissue types at early stages. Interestingly, activation of *Pax7-CreERT2* by injection a day earlier (at E8.5), resulted in a higher proportion of SOX9+, RFP+ cells in MCT in the deep pectoral girdle muscles, further confirming their developmental origin in dermomyotome. Further, the finding that iBAT and sBAT brown adipocyte progenitors are labelled by *Pax7-CreERT2* and SOX9 with the same timing to selected connective tissues (MCT, myotendinous junctions) of deep pectoral muscles suggests a common developmental origin (discussed below).

Thus, to visualize brown adipocyte precursors prior to depot formation, we injected tamoxifen at E9.5 in Pax7-CreERT2; RFP embryos, and embryos were allowed to develop to E11.5. At cervical levels (C4-C5), hypaxial and epaxial muscle blocks were lineage-labeled (**Figure S4A**) as well as a triangle of cells that included the dorsal dermis and extended ventro- medially (**Figure 4A, S4A,B).** At the ventromedial tip of this triangle, we observed a small SOX9- positive population co-expressing EBF2 (**Figure 4Ai-v, B; S4A**) but not GATA6 (**Figure S4Ei-iii**). These SOX9-expressing cells were situated at the position where the sBAT depot will form: the boundary between the medially located epaxial muscles and the laterally-located hypaxial muscles (the latter will give rise to the deep pectoral girdle musculature, including the rhomboid and levator scapula (Saberi et al., 2017) (**Figure 4B; S4Ai-iv**). This population of SOX9-positive cells was adjacent to and appeared continuous with the developing vertebral transverse process, which also expresses SOX9 but is not descended from *Pax7*-expressing cells, underscoring the utility of the *Pax7* lineage label (**Figure S4Ei, ii**, blue arrows).

**Figure 4.**
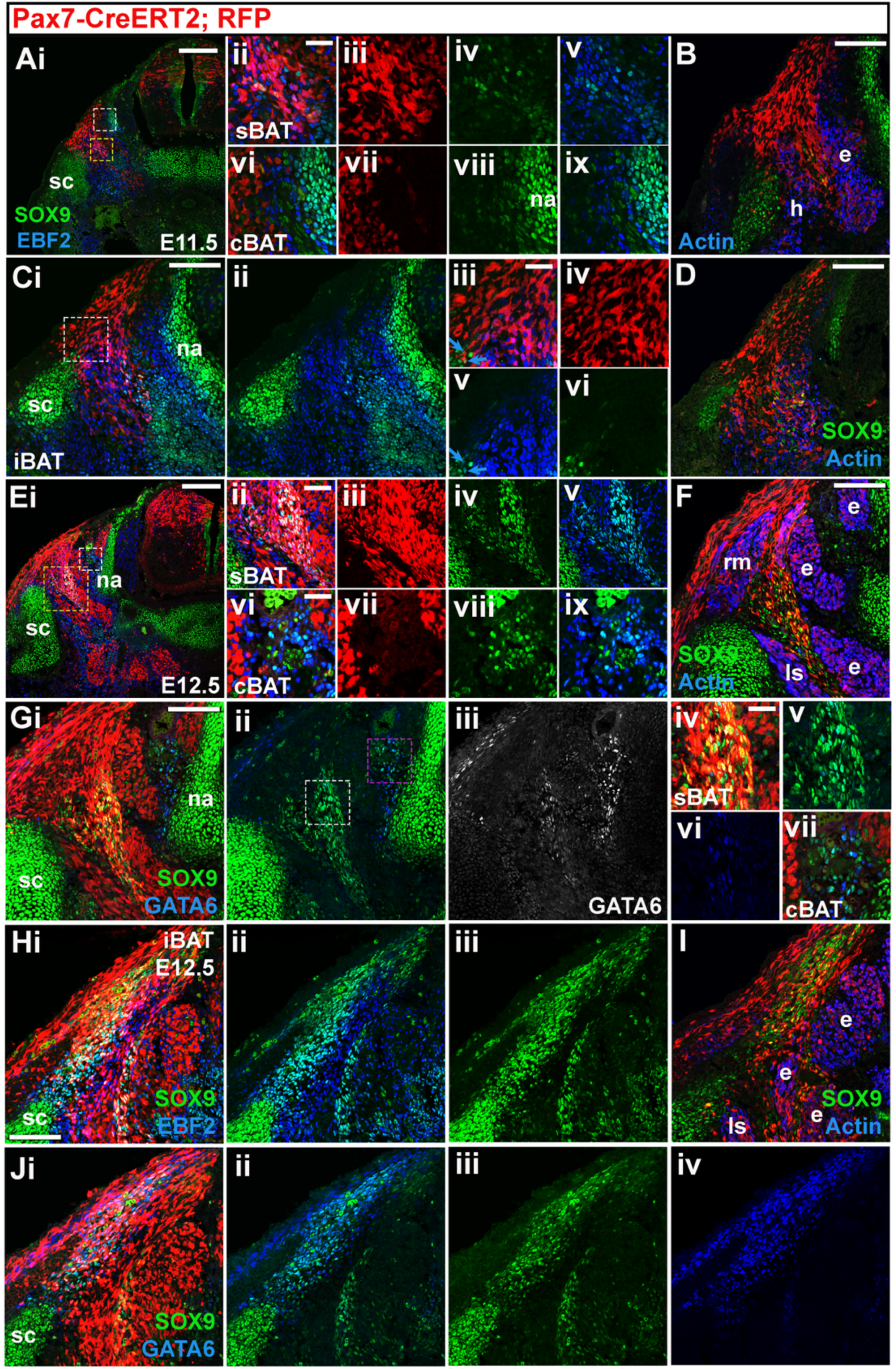
Spatial localization of SOX9-expressing brown adipocyte progenitors at early stages of development. (Ai) Immunofluorescence for SOX9, EBF2, and RFP (for the *Pax7* lineage) on cervical sections of *Pax7^Cre/ERT2/+^; Rosa26^tdTomato/+^* embryos injected with tamoxifen at E9.5 and examined at E11.5. Area of (Ai) bounded by yellow box is shown at higher magnification in (ii), and as RFP alone (iii), SOX9 alone (iv), and with SOX9 and EBF2 (v). Area of (Ai) bounded by white box is shown at higher magnification in (vi), and as RFP alone (vii), SOX9 alone (viii), and with SOX9 and EBF2 (ix). (B) Immunofluorescence for SOX9, Muscle Actin, and RFP in a similar section of a littermate to embryo shown in (A). (Ci) Immunofluorescence for SOX9, EBF2, and RFP, or SOX9 and EBF2 alone (ii) on anterior thoracic sections of *Pax7^Cre/ERT2/+^; Rosa26^tdTomato/+^* embryos injected with tamoxifen at E9.5 and imaged at E11.5. Area of (Ci) bounded by white box is shown at higher magnification in (iii), and as RFP alone (iv), EBF2 and SOX9 (v), and SOX9 alone (vi). Blue arrows in (Ciii) indicate SOX9-expressing *Pax7* descendants located near the dorsomedial border of the scapula. (D) Immunofluorescence for SOX9, Muscle Actin, and RFP in in a similar section of littermate to embryo shown in (C). (Ei) Immunofluorescence for SOX9, EBF2, and RFP on cervical sections of *Pax7^Cre/ERT2/+^; Rosa26^tdTomato/+^* embryos injected with tamoxifen at E9.5 and imaged at E12.5. Area of (Ai) bounded by yellow box is shown at higher magnification in (ii), and as RFP alone (iii), SOX9 alone (iv), and in SOX9 and EBF2 (v). Area of (Ai) bounded by white box is shown at higher magnification in (vi), and as RFP alone (vii), SOX9 alone (viii), and in SOX9 and EBF2 (ix). (F) Immunofluorescence for SOX9, muscle actin, and RFP in in similar section of littermate to embryo shown in (E). (G) Immunofluorescence for Sox9, GATA6, and RFP (i), SOX9 and GATA6 (ii), and GATA6 alone (iii), in an alternate section to that shown in (A). Area of (Gii) bounded by white box is shown at higher magnification in (v), including RFP (iv), and GATA6 alone (vi). Area of (Gii) bounded by pink box is shown at higher magnification in (vii) with RFP. (Hi) Immunofluorescence for SOX9, EBF2, and RFP, SOX9 and EBF2 (ii), and SOX9 alone (iii) on anterior thoracic sections of *Pax7^Cre/ERT2/+^; Rosa26^tdTomato/+^* embryos injected with tamoxifen at E9.5 and imaged at E12.5. (I) Immunofluorescence for SOX9, Muscle Actin, and RFP in in a similar section of littermate to embryo shown in (C). (Ji) Immunofluorescence for SOX9, GATA6, and RFP, SOX9 and GATA6 (ii), SOX9 alone (iii), and GATA6 alone (iv) in an alternate section to that shown in (H). Scale bars: 200µm: (Ai), (B), (Ci), (D), (Ei), (F), (Gi), (Hi); 50µm: (Eii); 25µm: (Aii), (Ciii), (Evi), (Giv). Abbreviations: cBAT, cervical BAT; e, epaxial muscle; h, hypaxial muscle; iBAT, interscapular BAT; na, neural arch; ls, levator scapula; na, neural arch; rm, rhomboid major; sBAT, scapular BAT; sc, scapula.

We identified a second population of SOX9-expressing cells in the E11.5 cervical region, located between the single epaxial muscle block and the neural arch (**Figure 4Ai, vi-ix**), a site consistent with the position of the cBAT depot. As would be predicted from our *Pax7* lineage analysis, these SOX9-expressing cells were not *Pax7* descendants (**Figure 3**; **Figure 4Ai, vi-vii**). We also detected EBF2-expressing cells in this region (**Figure 4Ai, ix**), but none expressing GATA6 (**Figure S4Ei-iii**). The EBF2+ cells appeared intermingled with the SOX9+ cells, suggesting that if either of these are cBAT precursors, their gene expression profile differs from that of the sBAT progenitors, which co-express SOX9 and EBF2 at E11.5 (**Figure 4Av**). Given that our *Sox9* lineage analysis suggested that a proportion of cBAT progenitors transiently down-regulate SOX9 expression at E11.5-E12.5 (**2Aii-Fii, S2)**, it is possible that both SOX9+ and EBF2+SOX9- cells are adipocyte progenitors. However, this also illustrates the difficulty of distinguishing SOX9+ adipocyte progenitors from other SOX9+ cell types without the aid of the *Pax7* lineage label.

Our *Sox9* lineage analysis suggested that SOX9-expressing iBAT progenitors within the *Pax7* descendant domain should be present in the anterior thoracic segments of E11.5 embryos. The iBAT depot is located medial to the scapula and superficial to the rhomboid muscles (**Figure S1E**). At E11.5, the hypaxially-derived rhomboid muscles were not yet in their dorsal position (**Figure S4Bi, iv**), but we found a few SOX9-expressing, *Pax7* descendant cells that were also co-expressing EBF2 medial to the scapula (**Figure 4Ci-vi; D, S4F**; blue arrows). These are candidate iBAT progenitors, and/or those of the medial scapular blade or deep pectoral girdle muscle connective tissues, given the restriction of the SOX9+ *Pax7* lineage observed above. We observed GATA6-expressing cells in this region as well, but they appeared distinct from the SOX9 population (**Figure S4Fi-iii**). Either (or both) of these populations may constitute iBAT adipocyte progenitors.

A day later, at E12.5, all three depot populations were identifiable in their mature positions relative to surrounding muscles. At cervical levels, the sBAT SOX9-expressing *Pax7* descendants formed a Y-shaped domain, located medial to the scapula and bounded on all sides by muscle--with the hypaxial levator scapulae positioned laterally to the sBAT, the epaxial muscles medially, and the rhomboid muscles dorsally (**Figure 4Ei, F; S4C**). Ventromedially, the sBAT domain at E12.5 could now be seen abutting the developing transverse process, which also expresses SOX9 and EBF2, as well as Tenascin (**Figure S4G)**, but is not descended from the *Pax7* expressing cells (**Figure S4Ei-ii**). SOX9 and EBF2 were co-expressed in the sBAT progenitors, and GATA6 expression was detectable as well (**Figure 4Ei-v, Gi-v**). The candidate cBAT progenitors, lying between the medial and intermediate epaxial muscle columns (**Figure 4Ei, vi-ix**), were more abundant at E12.5, and among the SOX9 positive cells in this region, we observed a population co-expressing EBF2 as well as GATA6 (**Figure 4Ei, vi-ix, Gi-iii, vii**). Additionally, the candidate iBAT progenitors, medial to the scapula and overlying the epaxial muscles, were now co-expressing SOX9, EBF2, and GATA6 (**Figure 4H-J; S4D; S1E**).

The *Sox9^CreERT2^* time course suggests that SOX9+ sBAT and iBAT adipocyte progenitors, labeled by E9.5 injection, may be present by E11.0. This coincides with the time when the central dermomyotome sheet that gives rise to them de-epithelializes. At E10.5, the dermomyotome is completely epithelial, as visualized by PDGFRα expression, which outlines cell membranes (**Figure S4H**). By E11.0, EMT has begun, and subset of de-epithelialized cells are *Pax7* descendants (**Figure S4Ii**), as visualized 18 hours after an E10.25 tamoxifen injection. In transverse sections, we were able to identify a small number of RFP+, SOX9+ cells lateral to the myotome (**Figure S4Ji-v**, blue arrows), and near the developing scapula (pink arrow), in a position where we would expect the earliest sBAT and iBAT progenitors to emerge from dermomyotome.

Taken together, our lineage and early developmental analyses demonstrate that SOX9 is a marker of early BAT progenitors, and have established a time-frame during which each of the three major depots form.

In order to further support the identification of SOX9+ cells as brown adipocyte progenitors, and to characterize their transcriptome, scRNA-Seq was performed on samples from E11.5-E13.5, a time spanning early detection of SOX9^+^EBF^+^ cells in de-epithelialized dermomyotome to when the three depots are morphologically distinct and lineage commitment, including expression of committed pre-adipocyte markers *Pparg* and *Cebpa* mRNAs (but not yet proteins), has occurred. Samples containing whole trunk tissue from the third cervical to second thoracic segments (C3-T2; a region containing the majority of all three BAT depots; **Figure S1**) were generated by dissection and dissociation to single cells. Limb and thoracic organ tissue was removed. After quality controls (see methods), a total of 23,299 cells were analyzed from 3 replicate samples at each of the 3 embryonic stages. Twenty-two clusters were generated at an initial Leiden resolution of 0.4, and cellular identity was determined using standard markers, following several recent analyses of similar embryonic samples (**Figure S5A-C)** (Fung et al., 2022; Jun et al., 2023; Rao et al., 2023).

Each of the tissue types we expected to recover based on the dissection strategy were observed. As shown in **Figure S5 Ci** and **Table S2**, clusters were obtained corresponding to neural progenitors and neural crest (*Sox2)*, differentiating neurons (*Elavl3)* including motor neurons (*Lhx1*), and Schwann cells (*Sox10, Mpz*). Additional clusters corresponded to epidermis (*Krt5*, *Krt14*), white blood cells (*Fcerg1, Tyrobp, Srgn, Csf1r*), and endothelium (*Cdh5, Kdr)* (**Figure S5Cii**). Three clusters expressed skeletal muscle markers including *MyoD1 and Pdgfa*; of these, one corresponded to differentiating myocytes (*Myh3, Myog*) and the others included muscle progenitors and dermomyotome, (*Myog*, *Pax7* and *Pax3*) **(Figure S5Ciii)**. Virtually all of the non-neural *Pax7*-expressing cells were contained within a dermomyotome cluster, consistent with previous findings also replicated in this study that the dermal and brown adipocyte linages extinguish *Pax7* expression at or prior to E11.5 (Lepper & Fan, 2010).

A central group of clusters comprised *Pdgfra+* cells (**Figure S5D**). *Pdgfra* is broadly expressed in most non-myogenic mesoderm, including the non-myotome compartments of somites, the lateral plate mesoderm, and their derivatives, and including uncommitted brown adipocyte progenitors and committed brown pre-adipocytes (Jun et al., 2023; Rao et al., 2023). Based on our dissection strategy, each of these cell types was expected to be recovered. The 0.4 Leiden resolution did not fully segregate *Pdgfra+* cells according to known cell types (**Figure S5B**); for example, MCT and dermis (*Osr2* and *Twist2*, respectively) were contained within a single cluster, as were cells expressing tendon and brown pre-adipocyte markers (*Scx, Tnmd* and *Cebpa, Pparg,* respectively) and smooth muscle and cartilage (*Acta2* and *Col2a1,* respectively). Therefore, *Pdgfra+* cells were analyzed at Leiden resolution of 0.6, which produced separate clusters of *Pdgfra+* cells for each expected committed cell types (**Figure 5A** and **Table S3**), including meninges (*Cldn11, Foxc1*), chondrocytes (*Col2a1*), muscle connective tissue (*Osr2,Crabp1*), smooth muscle (*Tagln, Acta2*) and two clusters each of dermis (*Twist2*), and tendon *(Scx, Tnmd)*. A single cluster differentially expressed pre-adipocyte markers *Pparg* and *Cebpa*. Each of these clusters, containing committed cell types, consisted primarily cells from the E13.5 sample, with some contribution from the E12.5 sample (compare **Figure 5A** and **C**). The remaining *Pdgfra+* clusters were composed of younger, E11.5-E12.5 cells, including those expressing markers for sclerotome (*Pax1, Pax9, Col2a1*), syndetome (*Scx*), de-epithelialized dermomyotome (*Twist2, En1* but not *Pax7*) (**Figure S5Diii-v**), and pan-fibroblast markers like *Fn1* and *Vim* (not shown), *Sox9* and *Ebf2* (**Figure S5Dvi**). Collectively, we termed these clusters fibroblast progenitors (**Figure 5A**) following terminology of a previous report (Rao et al., 2023). Three fibroblast progenitor clusters contained the majority of E11.5 cells, while the fourth was primarily derived from the E12.5 sample. While the Leiden clustering underscores the transcriptomic similarity of E11.5-E12.5 fibroblast progenitors from diverse sources, there is also clear spatial segregation on the UMAP projection of cells that express markers of different mesodermal populations such as cartilage/sclerotome (*Col2a1, Pax1, Pax9*) or adipocytes (*Ebf2, Gata6, Cdh4)* consistent with the fact that lineage divergence has already occurred (**Figure S5C**). However, few E11.5-E12.5 cells expressed pre-adipocyte markers or clustered with brown pre-adipocytes, as expected given the late expression of adipocyte markers.

**Figure 5.**
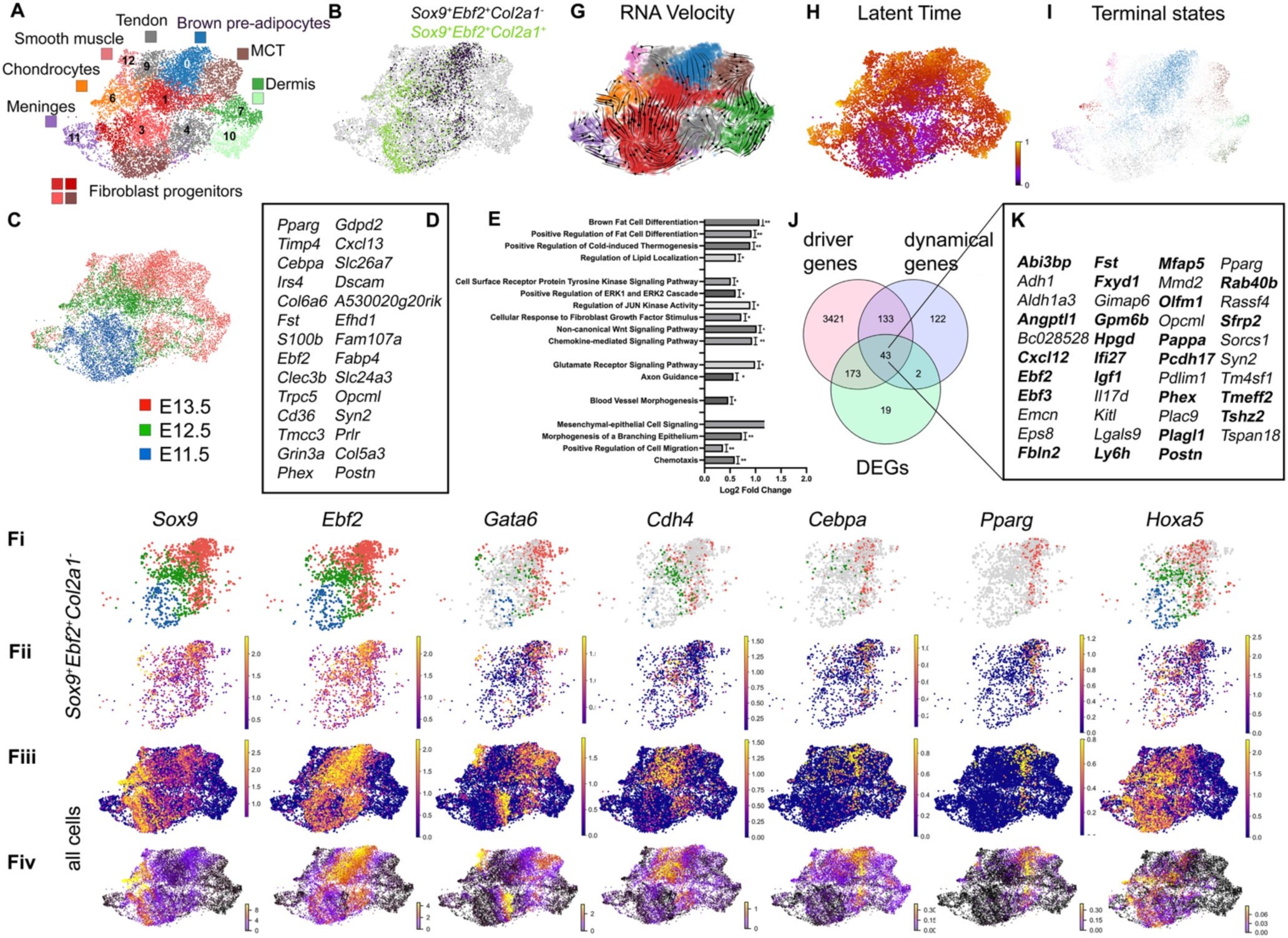
scRNA-Seq analysis to identify the brown pre-adipocyte lineage and trajectory (A-C) UMAP projection of *Pdgfra+* clusters at Leiden 0.6 resolution and labelled for cluster identity based on known markers (A), distribution of *Sox9+Ebf2+* cells subsetted based on *Col2a1* expression (B) or sample stage (C). (D) Top 20 genes differentially expressed in *Sox9+Ebf2+Col2a1-* cells compared to all other cells in the *Pdgfra+* clusters, sorted by lowest p-adjusted value. (E) Selected GO-Biological Process terms statistially over-represented among the genes differentially expressed in the same comparison. (F) Heat maps for key brown adipocyte progenitor and brown pre-adipocyte marker genes shows the sample of origin and spatial distribution on the UMAP projection. (Fi-ii) Show only *Sox9+Ebf2+Col2a1-* subestted cells color-coded by stage (i) or expression of the indicated brown adipocyte markers (ii). (Fiii-iv) heatmaps show expression of indicated markers in all *Pdgfra+ cells* with (iii) showing expression data based on exon data only and (iv) showing spliced/unspliced ratio from RNA velocity analysis. (G) RNA velocity vectors in *Pdgfra+* clusters. (H) Latent time of all cells in *Pdgfra+* clusters from RNA velocity analysis (I) Macrostates identified with CellRank2 by integration of sample stages and RNA velocity. (J) Comparison of top dynamical (RNA velocity) and driver (CellRank2) genes in the brown adipocyte trajectory to genes differentially expressed in the *Sox9+Ebf2+Col2a-* subset reveals substantial overlap. (K) 43 genes found in each list. Gene names marked in bold were also present in each of the 3 lists when list sizes were equalized to the top 237 genes each.

We next employed a subsetting approach, using *Sox9* and *Ebf2* as a novel marker combination, to enrich for BA progenitors and pre-adipocytes within the *Pdgfra+* clusters (**Figure 5B**). Based on our IF analysis, SOX9+EBF2+ cells are almost entirely restricted to brown adipocytes and progenitors, and to a subset of cartilage; therefore *Sox9+Ebf2+* cells were further subsetted into *Col2a1+* vs. *Col2a1-* populations (**Figure 5B**). The majority (90%) of *Sox9+Ebf2+Col2a1-* cells were found in the following groups: the brown pre-adipocyte cluster (41%), the fibroblast progenitor clusters (29%), and the two clusters neighboring brown pre- adipocytes identified as tendon (17%) or MCT (3%). Subsetted cells were derived from all three stages (**Figure 5Fi**). *Sox9+Ebf2+Col2a1-* cells and *Sox9+Ebf2+Col2a1+* cells, including those contained within the same fibroblast progenitor clusters, were spatially segregated in the UMAP projection, with the former located closer to the brown pre-adipocyte cluster and the latter closer to the chondrocyte cluster (**Figure 5B**). A comparison of differentially expressed genes in *Sox9+Ebf2+Col2a1-* compared to all other cells in the *Pdgfra+* clusters showed that known preadipocyte markers *Pparg, Cebpa* and *Fabp4* were among the most highly differentially expressed genes (DEGs) (**Figure 5D** and **Table S4**). Gene ontology (GO) terms over-represented among DEGs included terms for brown adipocyte (and other adipocyte) differentiation (**Figure 5E** and **Table S5).** Together, this strongly suggests that the subsetting approach isolated a population of cells, using a novel marker combination of *Sox9* and *Ebf2*, that includes both brown pre-adipocytes and brown adipocyte progenitors spanning E11.5-E13.5. This provides an opportunity to observe gene expression and the signaling environment experienced by these cell populations (**Figure 5D-E, Figure S5E**, and discussed below).

Within the *Sox9+Ebf2+Col2a1-* subset, the timing of expression of known marker genes was consistent with published work and our IF results: *Sox9* and *Ebf2* were observed earliest and were present at all stages (E11.5-E13.5) while by *Cdh4* and *Gata6* were present in the *Sox9+Ebf2+Col2a1-* population at E12.5-E13.5 and finally preadipocyte markers *Pparg and Cebpa* were expressed primarily at E13.5 (**Figure 5Ei-ii)**. *Hoxa5,* previously shown to be expressed in the earliest visualized depots and to regulate depot development (Holzman et al., 2021), was also broadly expressed including *Sox9+Ebf2+Col2a1-* cells at all three stages **(Figure 5Eii-iii)**. Given that scRNA-seq under-samples, leading to false negatives for *Sox9* and *Ebf2* gene expression, we also examined heatmaps of these marker genes within the entire *Pdgfra+* dataset (**Figure 5iii**), which showed a highly similar pattern to the subsetted *Sox9+Ebf2+Col2a1-* cells. Excitingly, in addition to expected markers, several genes were differentially expressed and GO terms significantly over-represented among differentially expressed genes in the *Sox9+Ebf2+Col2a1-* subset, that have not been previously associated with embryonic brown adipocyte or dept development (**Figure 5D-E**). These are discussed below, and open avenues for further study.

In a second approach, we used RNA velocity analysis (La Manno et al., 2018) to model the trajectory of cells toward commitment to a brown pre-adipocyte fate. As shown in **Figure 5G**, this revealed that vectors progressing toward the brown pre-adipocyte cluster originated primarily from the E11.5-E12.5 fibroblast progenitor clusters, as expected. Most notable for our analysis, vectors originating from E11.5 fibroblast progenitor clusters progressed toward both the brown adipocyte and chondrocyte clusters, with a spatial pattern that overlapped closely with the distribution *Sox9+Ebf2+Col2a1-* and *Sox9+Ebf2+Col2a1+* subsets, respectively (compare **Figure 5G** to **B**). A latent time analysis (**Figure 5H**) of RNA velocity data was consistent with each of the committed (predicted oldest) cell types identified by clustering. Phase portraits of key brown adipocyte progenitor and brown pre-adipocyte markers indicated that the model well-captured the velocity of each gene (**Figure S5F**). The top dynamical genes associated with the brown pre-adipocyte trajectory are given in **Table S6** and, like DEGs from the subsetting approach, contained numerous known markers for pre-adipocytes, as well as additional genes whose role in depot formation or brown pre-adipocyte commitment have not yet been studied.

Finally, we applied CellRank2 (Weiler et al., 2024), an approach that integrates biological stage (embryonic day of each sample) with RNA velocity data to model cell trajectories. This approach identified terminal states that overlapped with the mature cell type identities assigned by Leiden clustering and marker gene analysis (compare **Figure 5I** to **A**). All of the Leiden clusters assigned to a mature cell type (**Figure 5A**) had a corresponding terminal state predicted by CellRank2 (**Figure 5I**), with the exception of the tendon cluster neighboring the brown adipocyte cluster; these two were identified by CellRank2 as a single terminal state.

Further, the heatmap showing trajectory of cells toward the brown pre-adipocyte terminal state overlaps considerably with the *Sox9+Ebf2+Col2a1-* subset described above; this convergence of results obtained with independent methods further strengthens the hypothesis that these cells represent brown pre-adipocyte progenitors as well as committed brown adipocytes, present in all three stages sampled. Driver genes associated with the brown pre- adipocyte trajectory, identified by CellRank2, are given in **Table S7.** Dynamical genes (from RNA velocity analysis) and driver genes (from CellRank2), are those whose transcription best fit the modeled trajectories toward the brown pre-adipocyte state, and are not independent since CellRank2 incorporates velocity data as one input. Notably, these three lists of genes associated with brown pre-adipocyte trajectory or identity showed considerable overlap and included both known markers of brown adipocyte fate (*Pparg, Ebf2*) and genes with unknown roles in BAT development *Sox9+Ebf2+Col2a1-* (**Figure 5J-K**). Finally, because the three gene lists were different in length, an additional comparison was made where each list was reduced to the top 237 genes; this comparison retained most of the same overlapping group of genes (indicated in bold in **Figure 5K**).

## Discussion

### SOX9 as part of a novel combinatorial marker for brown adipocyte progenitors and brown pre-adipocytes *in vivo*

SOX9 maintains numerous cell types in a proliferative, undifferentiated state (reviewed in (Lefebvre et al., 2007)). It is known to perform this function in white pre-adipocytes, where it directly represses pan-adipocyte transcriptional regulators *Cebpb* and *Cebpd*. *Sox9* loss of function accelerated white pre-adipocyte differentiation and conversely, forced expression prevented differentiation of white pre-adipocytes derived from adult mouse WAT (Gulyaeva et al., 2018; Y. Wang & Sul, 2009).

We find that SOX9 similarly marks brown adipocyte progenitors and pre-adipocytes during embryonic BAT depot formation *in vivo*, and that its expression precedes that of the mRNA or protein for committed brown pre-adipocyte markers such as *Cepba* and *Pparg*. Further, SOX9 is co-expressed with EBF2, a marker selective for brown, as opposed to white, brown adipocyte progenitors (J. Wang & Tontonoz, 2017; W. Wang et al., 2014) in sBAT and iBAT progenitors by E11.5. While EBF2 is broadly expressed in numerous mesoderm derivatives, the combination of SOX9 and EBF2 distinguishes brown adipocyte progenitors and pre-adipocytes from other cell types. By E12.5, SOX9 is also co-expressed with GATA6, an established marker of brown adipocyte progenitors (Jun et al., 2023; Rao et al., 2023). In the developing cBAT, the initial expression of SOX9, EBF2, and GATA6 was more dynamic, but by E13.5, all three markers were co-expressed. SOX9 is downregulated concomitant with the onset of PPARγ protein expression brown pre-adipoctyes of all depots, but expression persists in the PDGFRα-positive BAT connective tissue of each depot. SOX9’s expression pattern, used as part of a combinatorial marker, afforded the opportunity to observe brown adipocyte progenitors and brown pre-adipocytes at early stages.

Lineage-labeling with *Sox9-CreERT2* was used to assess the timing of *Sox9* activation in brown adipocyte progenitors and summarized in **Figure S2M.** The Sox9 lineage label was mostly absent from skeletal muscle and rare in dermis, which is important for its use to distinguish brown adipocyte progenitors from other dermomyotome derivatives. A few dermal fibroblasts were labelled by *Sox9-CreERT2* following tamoxifen injection at or before E10.5. However, dermal fibroblasts do not express EBF2 and are spatially restricted to a lateral position in contrast to the SOX9+EBF2+ adipocyte progenitors observed at E11.5, allowing the two to be distinguished. To further distinguish adipocyte progenitors from other SOX9+ cell types, including cartilage, tendon and MCT, we used the *Pax7-CreERT2* lineage label to mark descendants of central dermomyotome. iBAT and sBAT depots were fully labeled by tamoxifen injection at E8.5-E9.5, while other SOX9+ cell types including cartilage, tendon, and epaxial muscle MCT, were not labelled. Thus, the combination of SOX9 with EBF2 and/or *Pax7* lineage is sufficient to distinguish brown adipocyte progenitors and brown pre-adipoctyes of the iBAT and sBAT. Importantly, an exception is connective tissues of the deep pectoral girdle muscles, which were labelled under the same conditions that brown adipocytes were labelled (discussed further below).

Finally, using *Sox9* lineage labeling, we showed that cells labelled by E11.5 Tamoxifen injection (and thus expressing *Sox9* as early as E11.75-E12.5) also give rise to most adipocytes of the perinatal iBAT and sBAT depots, despite substantial growth of the depots during the fetal period (between ∼E14.5-E18.5). A similar density of labeling of the perinatal depots resulted when tamoxifen was delivered early at E11.5 or later at E15.5, a time when SOX9 expression is restricted to BAT connective tissue. This establishes SOX9+ PDGFRα+ connective tissue as a major source of new adipocytes during fetal expansion of the iBAT and sBAT depots, and suggests that the cells lineage labelled for *Sox9* early include progenitors of both brown adipocytes and BAT connective tissue.

### Developmental origin and embryonic position of the major BAT depots

Our finding that iBAT is fully labelled by *Pax7-CreERT2* and *Meox1-Cre* is consistent with previous reports (Lepper & Fan, 2010; Sebo et al., 2018), and the timing of *Pax7-CreERT2* labeling is nearly identical to the previous report (Lepper & Fan, 2010), despite using a different *Pax7Cre-ERT2* allele. All of this strongly supports the conclusion that iBAT is derived from central dermomyotome of somites.

Additionally, we show here that the sBAT and cBAT depots are entirely somite-derived and that the sBAT emerges from *Pax7+* dermomyotome with similar timing to iBAT. However, cBAT does not arise from *Pax7+* dermomyotome, leaving open its origin. Some cBAT has an expression history of *Pax3* (Sanchez-Gurmaches & Guertin, 2014), which could indicate an origin in the Pax*7-*negative dorsal (epaxial) or ventral dermomyotome; an origin in epaxial dermomyotome is consistent with its position enclosed by epaxial muscles (Buckingham & Relaix, 2015). However, cBAT only partially derives from *Pax3* positive cells. Given that we find cBAT adipocytes are also lineage-labeled by *Sox9Cre-ERT2* following E8.5 and E9.5 tamoxifen injection, a time that targets compartmentalized somites, a partial origin in sclerotome is also possible. Sclerotome, unlike dermomyotome, is highly SOX9+ in compartmentalized somites. A sclerotome-specific lineage labeling strategy could test this hypothesis in the future.

While the combination of SOX9 expression and *Pax7* lineage labeling distinguished iBAT and sBAT brown adipocytes from nearly all other cell types, our labeling strategy was unable to separate them from a population of cells contributing to the MCT and myotendinous junctions of deep pectoral girdle muscles, the rhomboid and levator scapula. This suggests that both populations arise with similar timing from the Pax7+ central dermomyotome and thus could derive from the same progenitor population and/or be co-regulated. The rhomboid and levator scapula are derived from hypaxial myotomes (Saberi et al., 2017). Their development differs from other muscles of the forelimb and pectoral girdle in at least two ways: the deep pectoral girdle muscles are primaxial: their MCT does not arise from lateral plate mesoderm (Durland et al., 2008), and they develop by a mode that does not involve muscle progenitor migration (Valasek et al., 2011). While a somitic origin for the deep pectoral girdle musculature is expected, to our knowledge, a dermomyotome origin, indicated by *Pax7* lineage labeling, has not been previously described for MCT. The development of the deep pectoral girdle musculature including its connective tissues is understudied, particularly the earliest aspects of its formation. It is notable that by scRNA-seq analysis, brown pre-adipocyte and MCT clusters were adjacent in the UMAP projection indicating transcriptional similarity.

The iBAT and sBAT depots both arise between the epaxial muscles and hypaxial-derived deep pectoral girdles muscles, at axial levels that overlap with the forelimb field and cervical-thoracic transition. It will be valuable to track the specific segments that give rise to iBAT and sBAT adipocytes, to compare them to those that form the deep pectoral girdle muscles and/or identify shared regulatory mechanisms. *Hoxa5,* which affects BAT development *in vivo* also plays roles in patterning skeletal and muscular structures across the cervical to thoracic transition (Holzman et al., 2021). In contrast, cBAT is located between epaxial muscles derived from the medial and intermediate epaxial columns. While it is fully somite-derived, additional lineage labeling strategies are needed to determine its somitic compartment(s) of origin.

### Transcriptomic profile of brown adipocyte progenitors

*Sox9, Ebf2* and *Col2a1* provided a novel marker combination that permitted a subsetting approach, in which we compared gene expression of putative brown adipocyte progenitors and brown pre-adipocytes to other *Pdgfra+* clusters, which broadly comprise non-muscle mesodermal lineages derived from somites and LPM. Although scRNA-Seq under-samples sequences per cell, and is expected to produce to false negatives in our *Sox9+Ebf2+* subsetting approach, nevertheless, *Pparg* and *Cebpa* were the first and third on the list of differentially expressed genes in this subset (listed by p-adjusted value), and numerous other brown adipocyte markers (and GO terms) were statistically enriched, supporting the validity of the subsetting approach. Further, the position of *Sox9+Ebf2+Col2a1-* cells on the UMAP substantially overlapped with the brown pre-adipocyte trajectory modeled by RNA velocity analysis, an independent method. Finally, a semi-supervised approach (supervised only in that approximately 10 macrostates were set as optimal) identified *Pparg* and *Cepba*-expressing committed pre-adipocytes as one macrostate (stable cell type) within the *Pdgfra+* population, and the modeled trajectory leading to this macrostate also substantially overlapped on the UMAP with the *Sox9+Ebf2+Col2a1-* subset. While the trajectories predicted by the RNA velocity and macrostates analysis are not independent (the latter uses the former as one input), they do involve different computational methods, and are both independent of the subsetting approach. Thus, their convergence on overlapping sets of known and novel markers for the brown adipocyte lineage gave confidence in our identification of the brown adipocyte linage in the dataset.

In addition to known brown pre-adipocyte markers and GO pathways, each analysis of the scRNA-Seq data also yielded genes and GO terms that have not been previously characterized in the context of BAT depot development, and that provide new avenues for study. There was considerable overlap in the genes and pathways uncovered by each approach, although each is expected to provide related but not identical information.

Among pathways and genes that appeared associated with the brown adipocyte lineage in our analyses, several were known to be important for adult BAT maintenance and/or expansion following cold stress, or for browning of white adipocytes, suggesting shared mechanisms of postnatal and embryonic brown adipogenesis and BAT organogenesis. For example, chemokine signaling regulates macrophage infiltration and, in turn, sympathetic innervation of adult mouse BAT, both of which are necessary for expansion during cold stress (for example (D. Lee et al., 2024) and reviewed in (W. Wang & Seale, 2016)). Fst, a secreted inhibitor of TGFß signaling, promotes the browning of white adipocytes (Braga et al., 2014), a finding with potential therapeutic applications (Li et al., 2019). BMP7 signaling has been shown to be necessary for embryonic depot formation, and to promote brown adipogenesis in adults (Boon et al., 2013; Elsen et al., 2014; Tseng et al., 2008), but the role of BMP levels or inhibition, as might be regulated by Fst or other secreted inhibitors, has not been explored in a developmental context. Pathways over-represented among genes differentially expressed in the brown adipocyte lineage also included Fgf and other ERK1/2 transduced pathways and non-canonical Wnt signaling. Fgf and Wnt signals are well-characterized in the context of somite patterning and cell fate specification; for example, Wnt ligands from the epidermis promote dermal fates in dermomyotome progenitors (Atit et al., 2006b), and Fgf regulates muscle as well as adjacent syndetome fate (reviewed in (Brent & Tabin, 2002). Both canonical and non-canonical Wnt signaling has been implicated in adult BAT, and the Frizzled receptors and secreted frizzled-related inhibitor SFRP2 identified in our embryonic brown pre-adipocyte lineage are all known to act in both pathways (reviewed in (Martinez-Marin et al., 2025)).

Several pathways and genes involved in mesenchymal-to-epithelial crosstalk were also identified, with brown adipocyte progenitors, as expected, expressing mesenchymal genes. In the context of depot formation, the relevant epithelial population is unknown, but candidates are the tissues neighboring early brown adipocyte progenitors. Future work can take advantage of the ability to mark early brown adipocyte progenitors and test the tissues and signals required for their development.

Overall, our scRNA-seq results are complementary to and broadly agree with previous findings that numerous mesodermal cell types arise from a relatively homogenous population of fibroblast progenitors (discussed below) (Fung et al., 2022; Jun et al., 2023; Rao et al., 2023). However, our sample collection was designed to include all mesodermal derivatives of the trunk at the axial levels where depots form, while other approaches enriched for BAT using a dissection strategy or sorting strategy (dorsal tissue/iBAT dissection or *Pax7* lineage sorting).

Thus, our work provides a global comparison of fibroblast progenitors from embryonic trunks, and yields information complementary to that of the above sources. This should provide an additional useful resource for studying emergence of brown pre-adipocytes within the context of surrounding mesodermal, and especially somite-derived, tissues *in vivo*.

### Somite-derived fibroblast lineages and evolution of a novel cell type from somites

Using scRNA-seq across multiple time points, we identified a population of *Pdgfra*-expressing fibroblast progenitors that resolve into descendants of both the dermomyotome and sclerotome. This initial convergence in a common fibroblast progenitor profile, followed by molecular divergence into distinct lineages such as dermis and brown adipocytes for dermomyotome, and cartilage for sclerotome, suggests the possibility of a shared developmental trajectory between the dermomyotome and sclerotome compartments of the developing somite. Both compartments undergo epithelial-to-mesenchymal transition (EMT) to generate mesenchymal fibroblast progenitors that subsequently differentiate, based on their distinct genetic histories and environmental contexts. A key distinguishing feature of dermomyotome-derived fibroblast progenitors is that they arise from a *Pax7+* lineage, whereas sclerotome-derived fibroblast progenitors do not. Mesenchymal lineage specification is further shaped by local signaling cues, including Wnt/β-catenin for dermis, and BMP and Shh for sclerotome and cartilage patterning (reviewed in (Brent & Tabin, 2002)).

A shared molecular trajectory among fibroblast progenitors may reflect the evolutionary origin of the dermomyotome and sclerotome compartments of vertebrate somites. Supporting this idea, analysis of somite compartmentalization in *Amphioxus*—the best extant model of the ancestral chordate—reveals that the non-myotome portion of the somite forms a thin, continuous epithelium composed of a dermatome-like external cell layer (ECL) and a sclerotome-like ventral domain (Mansfield et al., 2015; Yong et al., 2021). Both regions contain *ColA-*expressing connective tissue progenitors, which are thought to contribute to extracellular matrix (ECM) layers that provide axial support. This could suggest that the entire non-myotome somite domain in the ancestral chordate functioned as a unified source of fibroblast-like cells, later partitioned in vertebrates into dermomyotome and sclerotome compartments that give rise to related progenitor populations subsequently directed along distinct developmental trajectories. Intriguingly, like the vertebrate dermomyotome, the lateral region of the *Amphioxus* somite has been shown to express *Amphi Pax3/7*, and to be dependent on BMP signaling (Yong et al., 2021).

In vertebrates, the dermomyotome is a transient epithelial structure that gives rise to the dorsal dermis and, through successive waves of cell migration, forms the underlying myotome. Lineage tracing, transplantation, and live imaging studies in chick have shown that the *Pax7*-expressing central epithelial dermomyotome contains bipotent precursors which, coincident with initiating an epithelial-to-mesenchymal transition (EMT), form a population of mesenchymal cells referred to as the intermediate domain (ID) and go on—via oriented cell divisions—to give rise either to the overlying dorsal dermis or to the underlying myotome (Ben-Yair et al., 2003; Ben-Yair & Kalcheim, 2005; Gros et al., 2005; Relaix et al., 2005). In contrast to the early postmitotic myocytes generated at the dermomyotome borders, progenitors emerging from the central dermomyotome maintain *Pax7* expression, remain mitotically active, and contribute to the myotome as proliferative myotomal precursors (PMPs) (Ben-Yair et al., 2003; Ben-Yair & Kalcheim, 2005; Gros et al., 2005). These progenitors retain a mesenchymal identity and do not express markers of terminal muscle differentiation. Over developmental time, *Pax7*+ progenitors persist in embryonic and fetal muscle and eventually localize beneath the basal lamina in a position characteristic of postnatal satellite cells (Ben-Yair & Kalcheim, 2005). In contrast, the same study found that differentiation toward dermis from the ID is associated with downregulation of *Pax7*. The presence of a bipotent population within the central epithelial, and then mesenchymal, dermomyotome suggests that asymmetric fate determination may be governed by local environmental cues and/or cell-cell interactions (Ben-Yair & Kalcheim, 2005; Gros et al., 2005).

While most aspects of somite formation, compartmentalization, and differentiation are conserved between chick and mouse (Christ et al., 2007), the evolutionary transition to the mammalian somite includes the origin of brown adipocytes as an additional cell fate arising from the dermomyotome. It is possible that, in mammals, changes in gene expression, such as initiation of SOX9 in dermomyotome derivatives, local signaling from surrounding tissues, and/or cell–cell interactions could have enabled the evolutionary origin of this additional lineage. The *in vivo* labeling described above allowed us to visualize the positions and pattern of BAT depot emergence in the mouse somite, and to track brown adipocyte lineage segregation to around the time of dermomyotome de-epithelialization. The timing and PAX7 expression history of the sBAT and iBAT depots suggest that their progenitors may represent an additional lineage derived from a population of cells analogous to the *Pax7+* ID population described in the chick somite. Our combinatorial system for marking BAT progenitors, combined with single-cell gene expression profiling, provides a framework to begin dissecting the gene regulatory networks and local signaling cues that enable BAT progenitors to emerge as a mammalian-specific somitic cell fate.

## Acknowledgements

This work was funded by a grant from the National Science Foundation, IOS-2019537. The Barnard College confocal microscope used in this study was funded by National Science Foundation grant DBI-1828264. We thank students enrolled in the 2021-2022 and 2023-2024 *Project Lab in Molecular Genetics* courses (Barnard College) for analysis of pilot scRNA-seq data, and Lucie Jeannotte, Jean Charron, and members of their labs (Université Laval) for discussion of these pilot analyses. We thank members of the 2024-2025 *Project Lab in Molecular Genetics* and *RNA-Seq Analysis* courses (Barnard College) for preliminary analysis and discussion of the presented scRNA-Seq data. We thank Juha Partanen (University of Helsinki) for the *Meox^Cre^* mouse line; all other mouse lines were obtained from The Jackson Laboratory. We thank Jeremy Dasen (New York University) for the HOXA5 antibody and Erin Bush, Florencia Velez-Cortes and Simoni Tiano at the Columbia Sulzburger Genome Center for sequencing and analysis of the scRNA-Seq data, including generation of a combined dataset. Work for this project completed at the Columbia Sulzburger Genome Center was funded in part through the NIH/NCI Cancer Center Support Grant P30CA013696 and used the Genomics and High Throughput Screening Shared Resource, as well as the National Center for Advancing Translational Sciences, National Institutes of Health, through Grant Number UL1TR001873. The content is solely the responsibility of the authors and does not necessarily represent the official views of the NIH.

## Supplementary Table and Figure legends

**Table S1.** Antibodies used in this study

**Table S2.** Differentially expressed genes marking each cluster in the full, combined scRNA-seq dataset at Leiden resolution 0.4. Gene lists were generated in Loupe Browser 8 using the function to compare each cluster to the entire dataset.

**Table S3.** Differentially expressed genes marking each cluster in the *Pdgfra+* clusters from the combined scRNA-seq dataset at Leiden resolution 0.6. Gene lists were generated in Loupe Browser 8 using the function to compare each cluster to the entire dataset.

**Table S4.** Differentially expressed genes in *Sox9+Ebf2+Col2a1-* subset compared to all cells in the *Pdgfra+* clusters from the combined scRNA-seq dataset at Leiden resolution 0.6. Gene lists were generated in Loupe Browser 8 using the function to compare the subset to the entire dataset.

**Table S5.** GO:BP terms significantly overrepresented among the genes differentially expressed in the *Sox9+Ebf2+Col2a1-* subset (Table S4), identified using the Panther DB search tool (REF).

**Table S6**. Top 300 dynamical genes that best model the trajectory toward brown pre- adipocytes, from RNA velocity analysis.

**Table S7**. Driver genes (q<.05) that best model the trajectory toward brown pre-adipocytes, from CellRank2 analysis.

**Figure S1.**
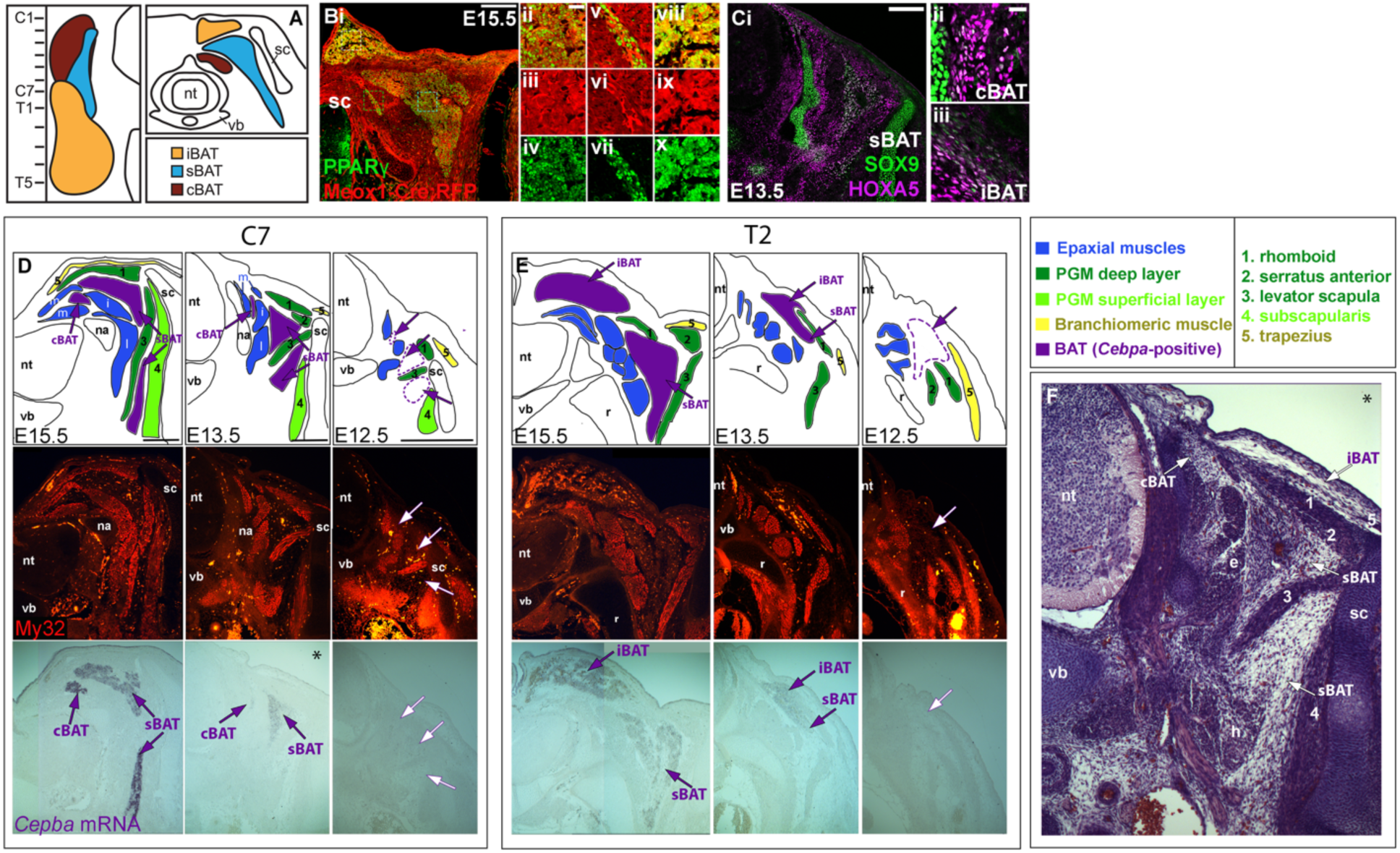
Identifying the locations and origins of developing BAT depots. (A) Schematic showing the location of mature BAT depots shown in dorsal (left) or transverse (right) views. In the left panel, anterior is up, the midline to the right with vertebrae labelled; the lateral is to the right. Redrawn from (Holzman et al., 2021). (B) Transverse section through *Meox1-Cre; RFP* embryo showing that all three depots are entirely labelled, and thus are all derived from somites. Magnified images are shown of the sBAT (ii-iv), cBAT (v-vii) and iBAT (viii-x) depots. (C) HOXA5 and SOX9 are co-expressed in brown adipocyte progenitors of all depots at E13.5 (iBAT is not shown). (D-E) *Cebpa* mRNA in situ hybridization (bottom row) and muscle actin immunofluorescence (middle row) were performed at the indicated stages, and the anatomical positions of muscles and BAT depots were schematized from these images (top row). Muscle identities are labelled as indicated in the legend (F) H&E stain of a tissue section adjacent to that shown in the middle column of panel (D). Scale bars: 200µm: (Bi, Ci); 25µm: Bii-x); 10µm: (Cii-iii): 250µm (D). Abbreviations: cBAT, cervical BAT; iBAT, interscapular BAT; sBAT, scapular BAT; na, neural arch; nt, neural tube; PGM, pectoral girdle muscles; r, rib; sc, scapula; vb, vertebral body. C1-C7, cervical vertebrae 1-7; T1-T5, thoracic vertebrae 1-5.

**Figure S2.**
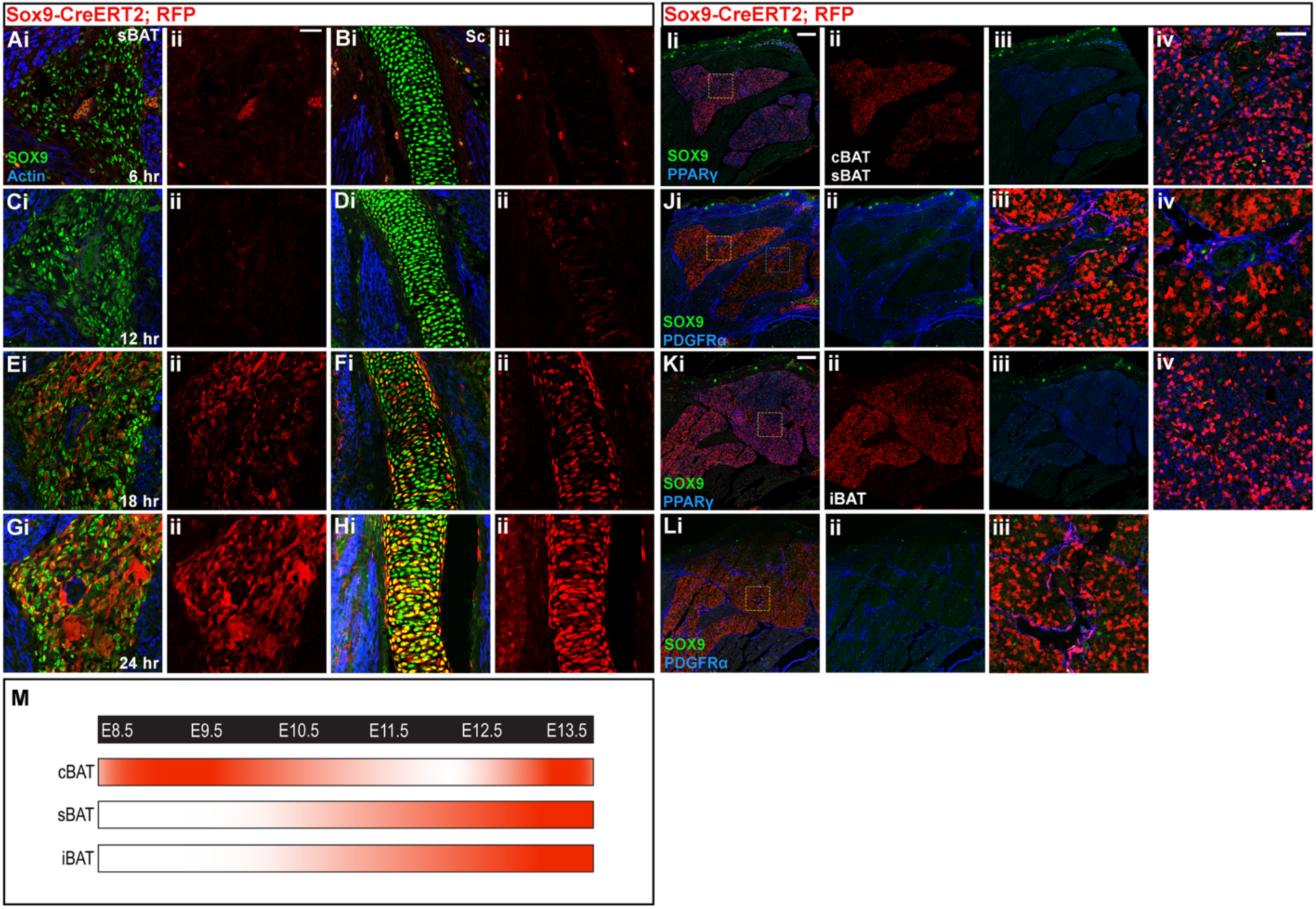
Characterization of Sox9-Cre activity and contribution of SOX9-expressing connective tissue to new brown adipocytes during fetal growth. (A-H) Timing of Sox9-Cre activation following a single injection of tamoxifen at E13.0 or E13.5 *Sox9^Cre/ERT2/+^; Rosa26^tdTomato/+^* embryos, times when *Sox9* is continuously expressed in brown adipocyte progenitors. Embryos were collected at 6 hours (A, B), 12 hours (C, D), 18 hours (E, F), or 24 hours (G, H) after injection and were examined for the presence of RFP+ cells. RFP+ cells are shown in the sBAT (A, C, E, G) and scapula (B, D, F, H). Red signal in (A) is red blood cell (RBC) autofluorescence. (I, K) Co-expression of PPARγ, SOX9, and RFP (for *Sox9* lineage) (i), RFP alone (ii), or PPARγ and SOX9 alone (iii), in transverse sections through either cervical (sBAT and cBAT) or thoracic (iBAT) regions in E18.5 *Sox9^Cre/ERT2/+^; Rosa26^tdTomato/+^* embryos. A single injection of tamoxifen at E15.5 was administered to examine the contribution of SOX9-expressing BAT connective tissue cells perinatal BAT and the end of the fetal growth period. (iv) Higher magnification view of the region indicated by the yellow box in (i) (sBAT). (J, L) Co-expression of PDGFRα (for connective tissue), SOX9, with (i) or without (ii) RFP (for *Sox9* lineage) in transverse sections through either cervical (sBAT and cBAT) or thoracic (iBAT) regions in alternate sections of those shown in (I, J). (iii) Higher magnification view of the region indicated by the yellow box in (i) (sBAT). (iv) Higher magnification view of the region indicated by the blue box in (i) (cBAT). (M) Schematic summarizing the time course of SOX9 expression in BAT progenitors of each depot, based on the results shown in Figure 2. Scale bars: 200µm: (I, Ki-iii; J, Li, ii), 50µm: (A-H; I, Kiv; Jiii, iv; Liii). Abbreviations: cBAT, cervical BAT; iBAT, interscapular BAT; sBAT, scapular BAT

**Figure S3.**
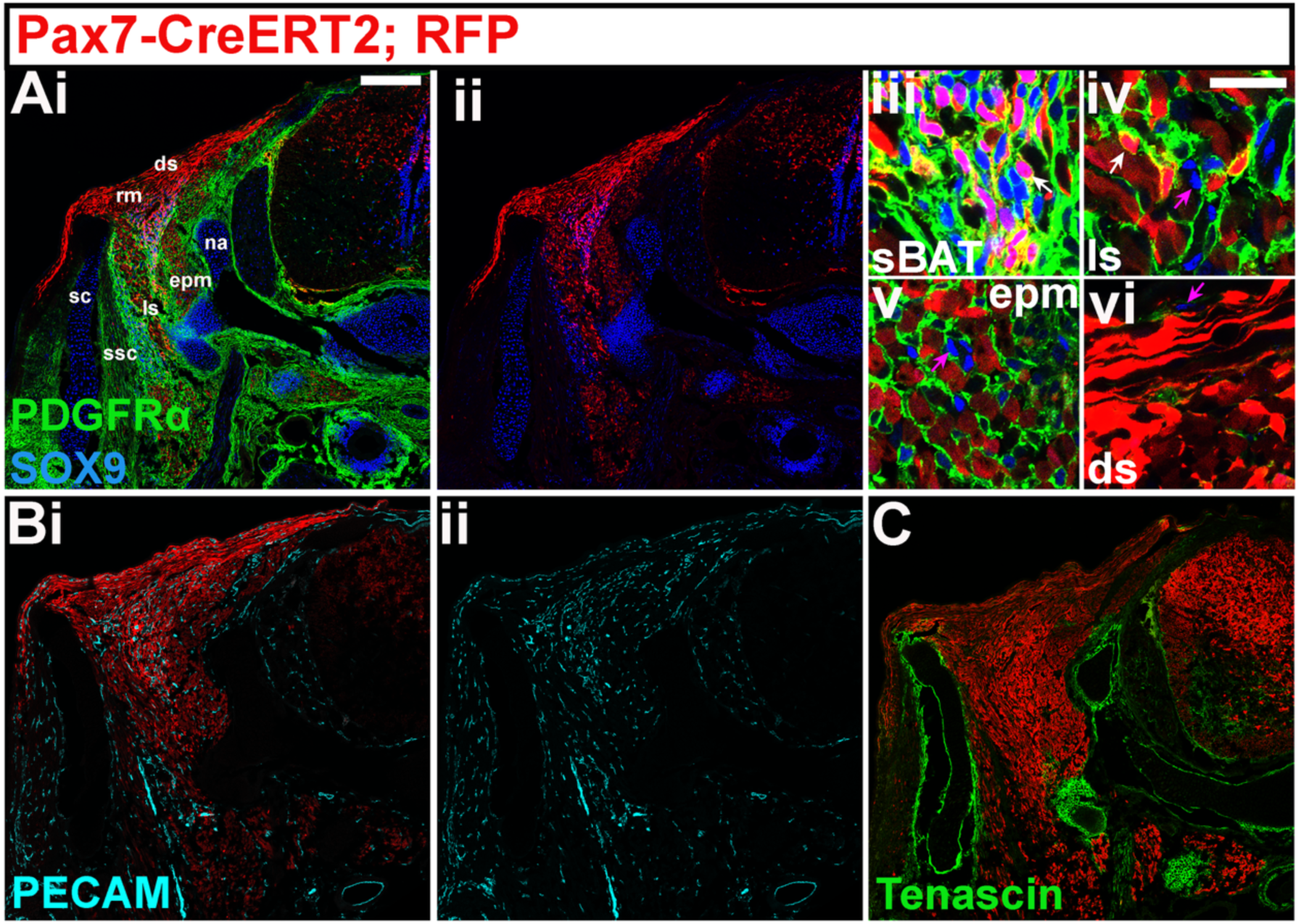
The majority of SOX9-expressing Pax7-descendants are found in BAT when Cre-ERT2 is activated by tamoxifen injection at E9.5. (Ai, ii) Immunofluorescence for SOX9, RFP (for the *Pax7* lineage), PDGFRα (ii) in cervical sections of *Pax7^Cre/ERT2/+^; Rosa26^tdTomato/+^* embryos injected with tamoxifen at E9.5 and examined at E14.5. Higher magnification views of (Ai) shown for (iii) sBAT, (iv) ls MCT, (v) ssc MCT, (vi) ds. Arrows in (Aiii-vi) mark the following: (iii) white arrow: SOX9+, RFP+ BAT connective tissue; (iv) white arrow: SOX9+, RFP+ MCT, pink arrow: SOX9+, RFP- MCT; (v) pink arrow: SOX9+, RFP- MCT; (vi) pink arrow: SOX9+, RFP- cell in dermis. (Bi, ii) Immunofluorescence for PECAM and RFP (for the *Pax7* lineage) in cervical sections of *Pax7^Cre/ERT2/+^; Rosa26^tdTomato/+^* embryos injected with tamoxifen at E9.5 and examined at E14.5. (C) Immunofluorescence for Tenascin and RFP (for the *Pax7* lineage) in cervical sections of *Pax7^Cre/ERT2/+^; Rosa26^tdTomato/+^* embryos injected with tamoxifen at E9.5 and examined at E14.5. Scale bars: 200µm: (Ai, ii), (B), (C); 25µm: (Aiii-vi). Abbreviations: ds, dermis; epm, epaxial muscles; ls, levator scapula; MCT, muscle connective tissue; na, neural arch; rm, rhomboid major; sc, scapula; ssc, subscapularis muscle.

**Figure S4.**
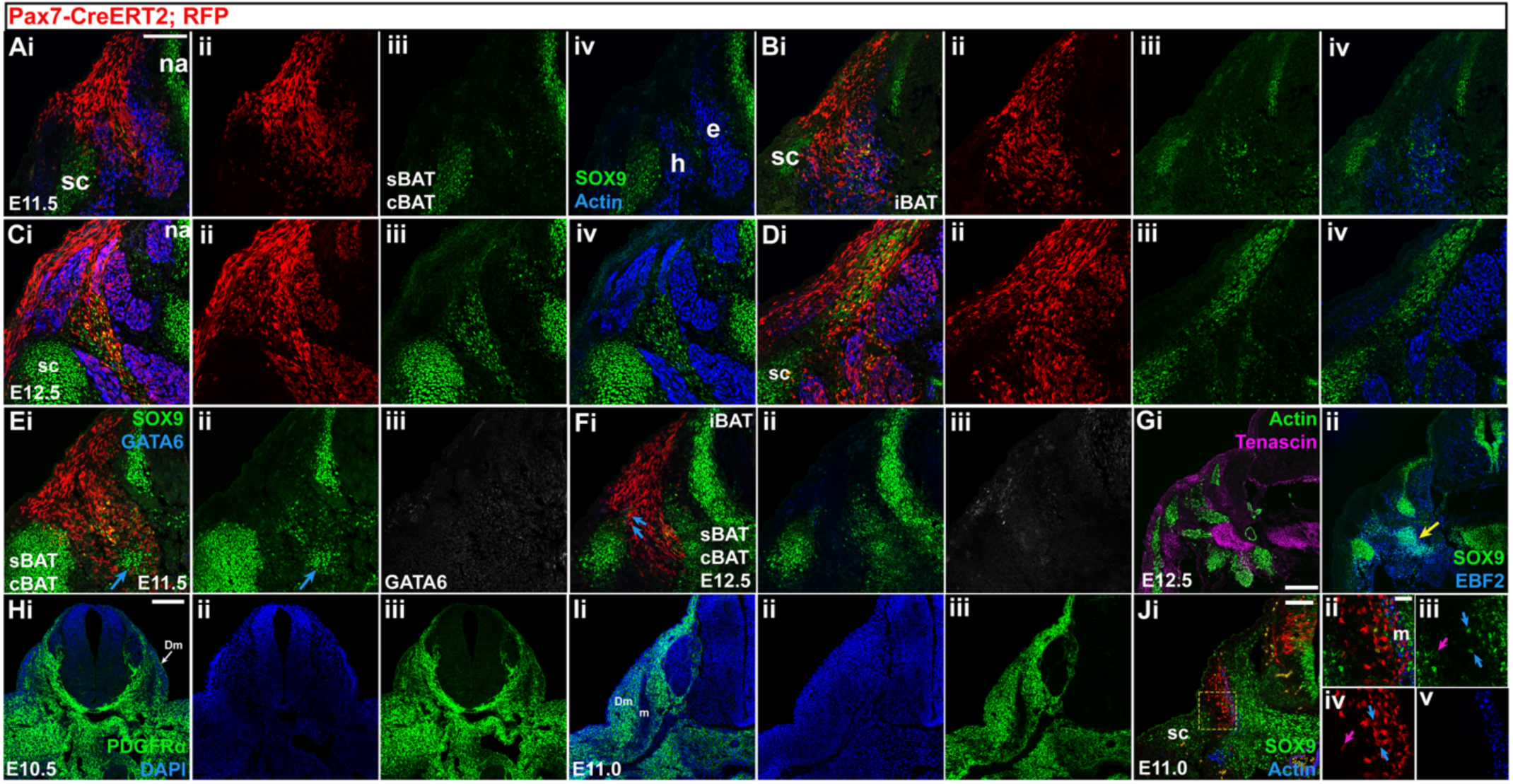
(Ai) Immunofluorescence for SOX9, muscle actin, and RFP (for the *Pax7* lineage), RFP alone (ii), SOX9 alone (iii), and SOX9 and muscle actin (iv), on cervical sections of E11.5 embryos from *Pax7^Cre/ERT2/+^; Rosa26^tdTomato/+^* mice injected with tamoxifen at E9.5. (B) Immunofluorescence for SOX9, muscle actin, and RFP, RFP alone (ii), SOX9 alone (iii), and SOX9 and muscle actin (iv), on anterior thoracic sections of E11.5 embryos from *Pax7^Cre/ERT2/+^; Rosa26^tdTomato/+^* mice injected with tamoxifen at E9.5. (C) Immunofluorescence for SOX9, muscle actin, and RFP, RFP alone (ii), SOX9 alone (iii), and SOX9 and muscle actin (iv), on cervical sections of E12.5 embryos from *Pax7^Cre/ERT2/+^; Rosa26^tdTomato/+^* mice injected with tamoxifen at E9.5. (D) Immunofluorescence for SOX9, muscle actin, and RFP, RFP alone (ii), SOX9 alone (iii), and SOX9 and muscle actin (iv), on anterior thoracic sections of E12.5 embryos from *Pax7^Cre/ERT2/+^; Rosa26^tdTomato/+^* mice injected with tamoxifen at E9.5. (Ei) Immunofluorescence for SOX9, GATA6, and RFP, SOX9 alone (ii), and GATA6 alone (iii), on cervical sections of E11.5 embryos from *Pax7^Cre/ERT2/+^; Rosa26^tdTomato/+^* mice injected with tamoxifen at E9.5. Blue arrow in Ei, ii indicates the SOX9+, RFP-transverse process.(Fi) Immunofluorescence for SOX9, GATA6, and RFP, SOX9 alone (ii), and GATA6 alone (iii), on anterior thoracic sections of E11.5 embryos from *Pax7^Cre/ERT2/+^; Rosa26^tdTomato/+^* mice injected with tamoxifen at E9.5. Blue arrows in (Fi) indicate SOX9-expressing *Pax7* descendants located near the dorsomedial border of the scapula. (G) Immunofluorescence on alternate sections through E12.5 embryos for (i) actin and tenascin and (ii) SOX9 and EBF2. Yellow arrow in Gii indicates the SOX9+, EBF2+ transverse process. A comparison of immunofluorescence on cervical sections for PDGFRα counterstained with DAPI (Hi, Ii), DAPI alone (Hii, Iii), or PDGFRα alone (Hiii, Iiii), at E10.5 (H) and E11.0 (I) reveals the transition from epithelial dermomyotome at E10.5, to a mesenchymal population at E11.0. (Ji) Immunofluorescence on an alternate section to that shown in (I) for SOX9, muscle actin, and RFP (for the *Pax7* lineage) on cervical sections of E11.0 embryos from *Pax7^Cre/ERT2/+^; Rosa26^tdTomato/+^* mice injected with tamoxifen at E10.25. Higher magnification view of the region indicated by the yellow box in (i) is shown in (ii), as SOX9 alone (iii), RFP alone (iv), and muscle actin alone (v). Blue arrows in (iii, iv) indicate Sox9+, RFP+ cells residing in the mesenchymal dermomyotome adjacent to the myotome. Pink arrow in (iii, iv) indicates a SOX9+, RFP+ cell near the scapula. Scale bars: 200µm: (A-G); 100µm: (H-Ji); 50µm: (Jii-v). Abbreviations: dm, dermomyotome; m, myotome; na, neural arch; sc, scapula.

**Figure S5.**
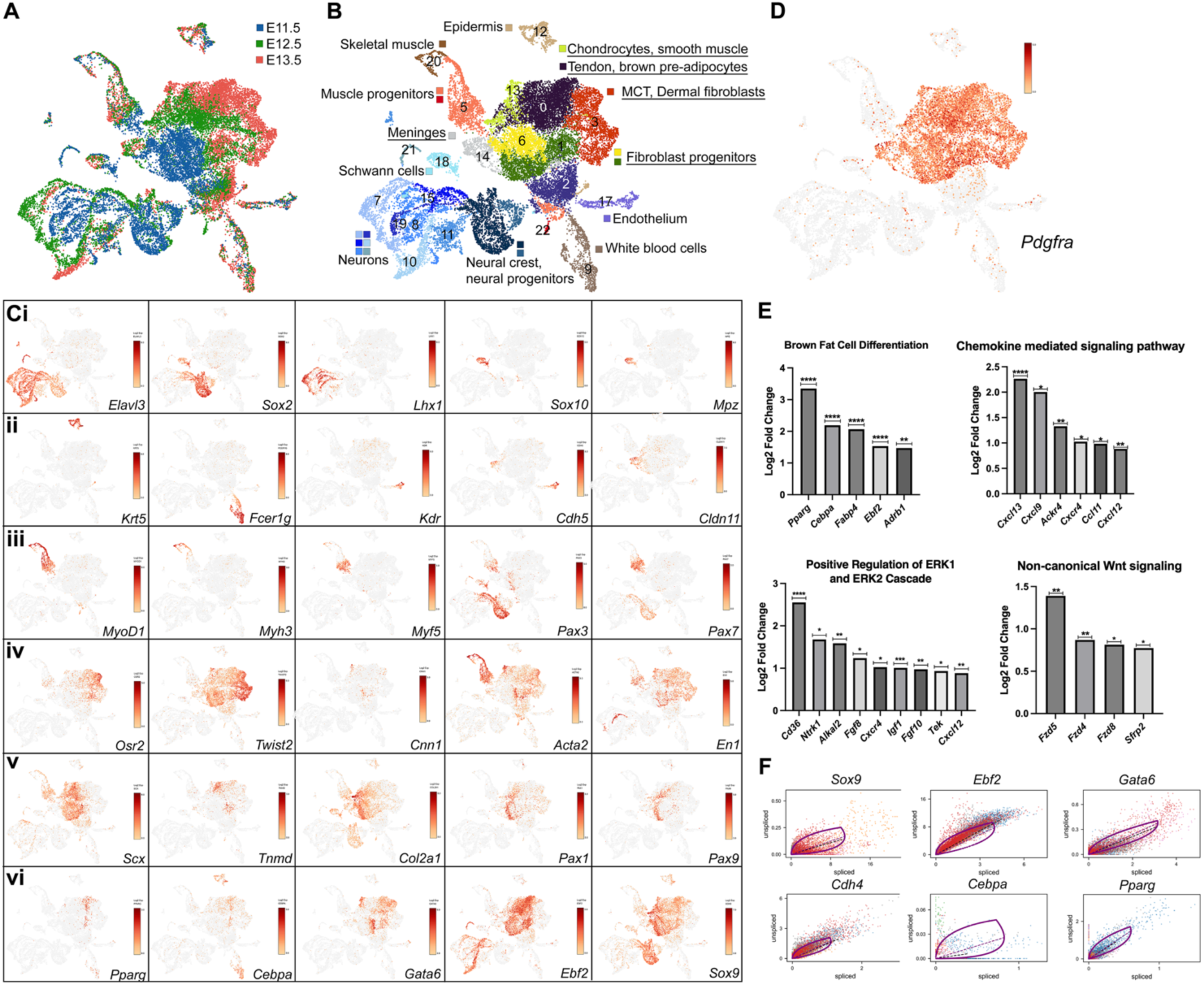
scRNA-seq transcriptional profiling of all cells from C3-T2 embryonic trunks, a population that includes brown adipoctye progenitors and brown pre-adipoctyes of the major depots (see text). UMAP projection of the E11.5-E13.5 combined sample is shown labelled for (A) sample of origin, and (B) Leiden clusters at resolution 0.4. Cell type identities contained within clusters are labelled, with the *Pdgfra+* clusters analyzed in Figure 5 underlined (see text). (C) Standard markers (see text) were used to assign cell type identities labelled in A, a subset of which are shown as feature maps, including markers for neurons and glia (Ci), epidermis, white blood cells, endothelium and meninges (Cii), skeletal muscle, muscle progenitors and dermomyotome (Ciii). (Civ-vi) show selected markers for cell types within the *Pdgfra+* clusters, including chondrocyte/smooth muscle, dermal fibroblast and meninges (Civ-v). (Cvi) shows brown pre-adipocyte and brown adipocyte progenitor markers including newly identified marker *Sox9*. (D) shows a feature map of *Pdgfra* expression. *Pdgfra+* clusters are those underlined in (B) and analyzed further in Figure 5. (E) Shows selected GO Biological Process pathways statistically over-represented in the *Sox9+Ebf2+Col2a1-* subset compared to all cells in *Pdgfra+* clusters, including each differentially expressed gene in our sample belonging to that pathway (* p-adj <0.05; ** p-adj <0.01; *** p-adj <.001; **** p-adj <.000). (F) Phase portraits of key markers of the brown adipocyte lineage show that the RNA velocity model well-caputres the velocity of each marker (see text and Figure 5).

## References

Ahmed, M. U., Maurya, A. K., Cheng, L., Jorge, E. C., Schubert, F. R., Maire, P., Basson, M. A., Ingham, P. W., & Dietrich, S. (2017). Engrailed controls epaxial-hypaxial muscle innervation and the establishment of vertebrate three-dimensional mobility. Developmental Biology, 430(1). 10.1016/j.ydbio.2017.08.011

Asou, Y., Nifuji, A., Tsuji, K., Shinomiya, K., Olson, E. N., Koopman, P., & Noda, M. (2002). Coordinated expression of scleraxis and Sox9 genes during embryonic development of tendons and cartilage. Journal of Orthopaedic Research, 20(4). 10.1016/S0736-0266(01)00169-3

Atit, R., Sgaier, S. K., Mohamed, O. A., Taketo, M. M., Dufort, D., Joyner, A. L., Niswander, L., & Conlon, R. A. (2006a). β-catenin activation is necessary and sufficient to specify the dorsal dermal fate in the mouse. Developmental Biology. 10.1016/j.ydbio.2006.04.449

Atit, R., Sgaier, S. K., Mohamed, O. A., Taketo, M. M., Dufort, D., Joyner, A. L., Niswander, L., & Conlon, R. A. (2006b). Β-Catenin Activation Is Necessary and Sufficient To Specify the Dorsal Dermal Fate in the Mouse. Developmental Biology, 296(1), 164–176. 10.1016/j.ydbio.2006.04.449

Ben-Yair, R., Kahane, N., & Kalcheim, C. (2003). Coherent development of dermomyotome and dermis from the entire mediolateral extent of the dorsal somite. Development, 130(18). 10.1242/dev.00667

Ben-Yair, R., & Kalcheim, C. (2005). Lineage analysis of the avian dermomyotome sheet reveals the existence of single cells with both dermal and muscle progenitor fates. Development, 132(4). 10.1242/dev.01617

Bergen, V., Lange, M., Peidli, S., Wolf, F. A., & Theis, F. J. (2020). Generalizing RNA velocity to transient cell states through dynamical modeling. Nature Biotechnology, 38(12). 10.1038/s41587-020-0591-3

Blondel, V. D., Guillaume, J. L., Hendrickx, J. M., De Kerchove, C., & Lambiotte, R. (2008). Local leaders in random networks. *Physical Review E - Statistical*, Nonlinear, and Soft Matter Physics, 77(3). 10.1103/PhysRevE.77.036114

Boon, M. R., van den Berg, S. A. A., Wang, Y., van den Bossche, J., Karkampouna, S., Bauwens, M., De Saint-Hubert, M., van der Horst, G., Vukicevic, S., de Winther, M. P. J., Havekes, L. M., Jukema, J. W., Tamsma, J. T., van der Pluijm, G., van Dijk, K. W., & Rensen, P. C. N. (2013). BMP7 Activates Brown Adipose Tissue and Reduces Diet-Induced Obesity Only at Subthermoneutrality. PLoS ONE, 8(9). 10.1371/journal.pone.0074083

Braga, M., Reddy, S. T., Vergnes, L., Pervin, S., Grijalva, V., Stout, D., David, J., Li, X., Tomasian, V., Reid, C. B., Norris, K. C., Devaskar, S. U., Reue, K., & Singh, R. (2014). Follistatin promotes adipocyte differentiation, browning, and energy metabolism. Journal of Lipid Research, 55(3). 10.1194/jlr.M039719

Brent, A. E., & Tabin, C. J. (2002). Developmental regulation of somite derivatives: Muscle, cartilage and tendon. Current Opinion in Genetics and Development, 12(5), 548–557. 10.1016/S0959-437X(02)00339-8

Buckingham, M., & Relaix, F. (2015). PAX3 and PAX7 as upstream regulators of myogenesis. In Seminars in Cell and Developmental Biology (Vol. 44). 10.1016/j.semcdb.2015.09.017

Cannon, B., & Nedergaard, J. (2004). Brown Adipose Tissue: Function and Physiological Significance. In Physiological Reviews (Vol. 84, Issue 1, pp. 277–359). Physiol Rev. 10.1152/physrev.00015.2003

Chen, J. W., Zahid, S., Shilts, M. H., Weaver, S. J., Leskowitz, R. M., Habbsa, S., Aronowitz, D., Rokins, K. P., Chang, Y., Pinnella, Z., Holloway, L., & Mansfield, J. H. (2013). Hoxa-5 acts in segmented somites to regulate cervical vertebral morphology. Mechanisms of Development, 130(4–5), 226–240. 10.1016/j.mod.2013.02.002

Christ, B., Huang, R., & Scaal, M. (2007). Amniote somite derivatives. Developmental Dynamics, 236(9), 2382–2396. 10.1002/dvdy.21189

Deries, M., Schweitzer, R., & Duxson, M. J. (2010). Developmental fate of the mammalian myotome. Developmental Dynamics, 239(11), 2898–2910. 10.1002/dvdy.22425

Durland, J. L., Sferlazzo, M., Logan, M., & Burke, A. C. (2008). Visualizing the lateral somitic frontier in the Prx1Cre transgenic mouse. Journal of Anatomy, 212(5), 590–602. 10.1111/j.1469-7580.2008.00879.x

Elsen, M., Raschke, S., Tennagels, N., Schwahn, U., Jelenik, T., Roden, M., Romacho, T., & Eckel, J. (2014). BMP4 and BMP7 induce the white-to-brown transition of primary human adipose stem cells. American Journal of Physiology - Cell Physiology, 306(5). 10.1152/ajpcell.00290.2013

Fung, C. W., Zhou, S., Zhu, H., Wei, X., Wu, Z., & Wu, A. R. (2022). Cell fate determining molecular switches and signaling pathways in Pax7-expressing somitic mesoderm. Cell Discovery 2022 8:1, 8(1), 1–21. 10.1038/s41421-022-00407-0

Gros, J., Manceau, M., Thomé, V., & Marcelle, C. (2005). A common somitic origin for embryonic muscle progenitors and satellite cells. Nature, 435(7044). 10.1038/nature03572

Gulyaeva, O., Nguyen, H., Sambeat, A., Heydari, K., & Sul, H. S. (2018). Sox9-Meis1 Inactivation Is Required for Adipogenesis, Advancing Pref-1+ to PDGFRα+ Cells. Cell Reports, 25(4). 10.1016/j.celrep.2018.09.086

Hachemi, I., & U-Din, M. (2023). Brown Adipose Tissue: Activation and Metabolism in Humans. Endocrinology and Metabolism, 38(2). 10.3803/EnM.2023.1659

Hayashi, S., & McMahon, A. P. (2002). Efficient recombination in diverse tissues by a tamoxifen-inducible form of Cre: A tool for temporally regulated gene activation/inactivation in the mouse. Developmental Biology, 244(2), 305–318. 10.1006/dbio.2002.0597

Holzman, M. A., Bergmann, J. M., Feldman, M., Landry-Truchon, K. I. M., Jeannotte, L., & Mansfield, J. H. (2018). HOXA5 protein expression and genetic fate mapping show lineage restriction in the developing musculoskeletal system. International Journal of Developmental Biology, 62(11–12), 785–796. 10.1387/ijdb.180214jm

Holzman, M. A., Ryckman, A., Finkelstein, T. M., Landry-Truchon, K., Schindler, K. A., Bergmann, J. M., Jeannotte, L., & Mansfield, J. H. (2021). HOXA5 Participates in Brown Adipose Tissue and Epaxial Skeletal Muscle Patterning and in Brown Adipocyte Differentiation. Frontiers in Cell and Developmental Biology, 9. 10.3389/fcell.2021.632303

Huang, R., Zhi, Q., Patel, K., Wilting, J., & Christ, B. (2000). Dual origin and segmental organisation of the avian scapula. Development, 127(17), 3789–3794. 10.1242/dev.127.17.3789

Jukkola, T., Trokovic, R., Maj, P., Lamberg, A., Mankoo, B., Pachnis, V., Savilahti, H., & Partanen, J. (2005). Meox1Cre: A mouse line expressing Cre recombinase in somitic mesoderm. Genesis, 43(3), 148–153. 10.1002/gene.20163

Jun, S., Angueira, A. R., Fein, E. C., Tan, J. M. E., Weller, A. H., Cheng, L., Batmanov, K., Ishibashi, J., Sakers, A. P., Stine, R. R., & Seale, P. (2023). Control of murine brown adipocyte development by GATA6. Developmental Cell, 58(21). 10.1016/j.devcel.2023.08.003

La Manno, G., Soldatov, R., Zeisel, A., Braun, E., Hochgerner, H., Petukhov, V., Lidschreiber, K., Kastriti, M. E., Lönnerberg, P., Furlan, A., Fan, J., Borm, L. E., Liu, Z., van Bruggen, D., Guo, J., He, X., Barker, R., Sundström, E., Castelo-Branco, G., … Kharchenko, P. V. (2018). RNA velocity of single cells. Nature, 560(7719). 10.1038/s41586-018-0414-6

Lee, D., Benvie, A. M., Steiner, B. M., Kolba, N. J., Ford, J. G., McCabe, S. M., Jiang, Y., & Berry, D. C. (2024). Smooth muscle cell-derived Cxcl12 directs macrophage accrual and sympathetic innervation to control thermogenic adipose tissue. Cell Reports, 43(5), 114169. 10.1016/j.celrep.2024.114169

Lee, Y. H., Petkova, A. P., Konkar, A. A., & Granneman, J. G. (2015). Cellular origins of cold-induced brown adipocytes in adult mice. FASEB Journal, 29(1). 10.1096/fj.14-263038

Lefebvre, V., Angelozzi, M., & Haseeb, A. (2019). SOX9 in cartilage development and disease. In Current Opinion in Cell Biology (Vol. 61). 10.1016/j.ceb.2019.07.008

Lefebvre, V., Dumitriu, B., Penzo-Méndez, A., Han, Y., & Pallavi, B. (2007). Control of cell fate and differentiation by Sry-related high-mobility-group box (Sox) transcription factors. In International Journal of Biochemistry and Cell Biology (Vol. 39, Issue 12). 10.1016/j.biocel.2007.05.019

Lepper, C., & Fan, C. M. (2010). Inducible lineage tracing of Pax7-descendant cells reveals embryonic origin of adult satellite cells. Genesis, 48(7), 424–436. 10.1002/dvg.20630

Li, H., Zhang, C., Liu, J., Xie, W., Xu, W., Liang, F., Huang, K., & He, X. (2019). Intraperitoneal administration of follistatin promotes adipocyte browning in high-fat diet-induced obese mice. PLoS ONE, 14(7). 10.1371/journal.pone.0220310

Madisen, L., Zwingman, T. A., Sunkin, S. M., Oh, S. W., Zariwala, H. A., Gu, H., Ng, L. L., Palmiter, R. D., Hawrylycz, M. J., Jones, A. R., Lein, E. S., & Zeng, H. (2010). A robust and high-throughput Cre reporting and characterization system for the whole mouse brain. Nature Neuroscience, 13(1), 133–140. 10.1038/nn.2467

Mansfield, J. H., Haller, E., Holland, N. D., & Brent, A. E. (2015). Development of somites and their derivatives in amphioxus, and implications for the evolution of vertebrate somites. EvoDevo, 6(1). 10.1186/s13227-015-0007-5

Martinez-Marin, D., Stroman, G. C., Fulton, C. J., & Pruitt, K. (2025). Frizzled receptors: gatekeepers of Wnt signaling in development and disease. Frontiers in Cell and Developmental Biology, 13, 1599355. 10.3389/fcell.2025.1599355

Mayeuf-Louchart, A., Lancel, S., Sebti, Y., Pourcet, B., Loyens, A., Delhaye, S., Duhem, C., Beauchamp, J., Ferri, L., Thorel, Q., Boulinguiez, A., Zecchin, M., Dubois-Chevalier, J., Eeckhoute, J., Vaughn, L. T., Roach, P. J., Dani, C., Pederson, B. A., Vincent, S. D., … Duez, H. (2019). Glycogen Dynamics Drives Lipid Droplet Biogenesis during Brown Adipocyte Differentiation. Cell Reports, 29(6), 1410–1418.e6. 10.1016/j.celrep.2019.09.073

McGlinn, E., Holzman, M. A., & Mansfield, J. H. (2019). Detection of Gene and Protein Expression in Mouse Embryos and Tissue Sections. *Methods in Molecular Biology (Clifton*, N.J.), 1920, 183–218. 10.1007/978-1-4939-9009-2_12

McInnes, L., Healy, J., Saul, N., & Großberger, L. (2018). UMAP: Uniform Manifold Approximation and Projection. Journal of Open Source Software, 3(29). 10.21105/joss.00861

Mekonen, H. K., Hikspoors, J. P. J. M., Mommen, G., Eleonore KÖhler, S., & Lamers, W. H. (2016). Development of the epaxial muscles in the human embryo. Clinical Anatomy, 29(8), 1031–1045. 10.1002/ca.22775

Mi, H., Muruganujan, A., Huang, X., Ebert, D., Mills, C., Guo, X., & Thomas, P. D. (2019). Protocol Update for large-scale genome and gene function analysis with the PANTHER classification system (v.14.0). Nature Protocols, *14*(3). 10.1038/s41596-019-0128-8

Murphy, M. M., Lawson, J. A., Mathew, S. J., Hutcheson, D. A., & Kardon, G. (2011). Satellite cells, connective tissue fibroblasts and their interactions are crucial for muscle regeneration. Development, 138(17). 10.1242/dev.064162

Oelkrug, R., Polymeropoulos, E. T., & Jastroch, M. (2015). Brown adipose tissue: physiological function and evolutionary significance. In Journal of Comparative Physiology B: Biochemical, Systemic, and Environmental Physiology (Vol. 185, Issue 6, pp. 587–606). Springer Verlag. 10.1007/s00360-015-0907-7

Rao, J., Djeffal, Y., Chal, J., Marchianò, F., Wang, C. H., Al Tanoury, Z., Gapon, S., Mayeuf-Louchart, A., Glass, I., Sefton, E. M., Habermann, B., Kardon, G., Watt, F. M., Tseng, Y. H., & Pourquié, O. (2023). Reconstructing human brown fat developmental trajectory in vitro. Developmental Cell, 58(21). 10.1016/j.devcel.2023.08.001

Relaix, F., Rocancourt, D., Mansouri, A., & Buckingham, M. (2005). A Pax3/Pax7-dependent population of skeletal muscle progenitor cells. Nature, 435(7044). 10.1038/nature03594

Saberi, M., Pu, Q., Valasek, P., Norizadeh-Abbariki, T., Patel, K., & Huang, R. (2017). The hypaxial origin of the epaxially located rhomboid muscles. Annals of Anatomy, 214, 15–20. 10.1016/j.aanat.2017.05.009

Sacks, H., & Symonds, M. E. (2013). Anatomical locations of human brown adipose tissue: Functional relevance and implications in obesity and type 2 diabetes. In Diabetes (Vol. 62, Issue 6). 10.2337/db12-1430

Sanchez-Gurmaches, J., & Guertin, D. A. (2014). Adipocytes arise from multiple lineages that are heterogeneously and dynamically distributed. Nature Communications, 5. 10.1038/ncomms5099

Seale, P., Bjork, B., Yang, W., Kajimura, S., Chin, S., Kuang, S., Scimè, A., Devarakonda, S., Conroe, H. M., Erdjument-Bromage, H., Tempst, P., Rudnicki, M. A., Beier, D. R., & Spiegelman, B. M. (2008). PRDM16 controls a brown fat/skeletal muscle switch. Nature, 454(7207), 961–967. 10.1038/nature07182

Sebo, Z. L., Jeffery, E., Holtrup, B., & Rodeheffer, M. S. (2018). A mesodermal fate map for adipose tissue. Development (Cambridge), 145(17). 10.1242/dev.166801

Sebo, Z. L., & Rodeheffer, M. S. (2019). Assembling the adipose organ: Adipocyte lineage segregation and adipogenesis in vivo. Development (Cambridge), 146(7). 10.1242/dev.172098

Sugimoto, Y., Takimoto, A., Akiyama, H., Kist, R., Scherer, G., Nakamura, T., Hiraki, Y., & Shukunami, C. (2012). Scx+/Scx9+ progenitors contribute to the establishment of the junction between cartilage and tendon/ligament. Development (Cambridge), 140(11), 2280–2288. 10.1242/dev.096354

Thomas, P. D., Ebert, D., Muruganujan, A., Mushayahama, T., Albou, L. P., & Mi, H. (2022). PANTHER: Making genome-scale phylogenetics accessible to all. In Protein Science (Vol. 31, Issue 1). 10.1002/pro.4218

Tseng, Y. H., Kokkotou, E., Schulz, T. J., Huang, T. L., Winnay, J. N., Taniguchi, C. M., Tran, T. T., Suzuki, R., Espinoza, D. O., Yamamoto, Y., Ahrens, M. J., Dudley, A. T., Norris, A. W., Kulkarni, R. N., & Kahn, C. R. (2008). New role of bone morphogenetic protein 7 in brown adipogenesis and energy expenditure. Nature, 454(7207), 1000–1004. 10.1038/nature07221

Valasek, P., Theis, S., DeLaurier, A., Hinits, Y., Luke, G. N., Otto, A. M., Minchin, J., He, L., Christ, B., Brooks, G., Sang, H., Evans, D. J., Logan, M., Huang, R., & Patel, K. (2011). Cellular and molecular investigations into the development of the pectoral girdle. Developmental Biology, 357(1), 108–116. 10.1016/j.ydbio.2011.06.031

Valasek, P., Theis, S., Krejci, E., Grim, M., Maina, F., Shwartz, Y., Otto, A., Huang, R., & Patel, K. (2010). Somitic origin of the medial border of the mammalian scapula and its homology to the avian scapula blade. Journal of Anatomy, 216(4), 482–488. 10.1111/j.1469-7580.2009.01200.x

van Marken Lichtenbelt, W. D., & Schrauwen, P. (2011). Implications of nonshivering thermogenesis for energy balance regulation in humans. American Journal of Physiology. Regulatory, Integrative and Comparative Physiology, 301(2). 10.1152/AJPREGU.00652.2010

Virshup, I., Rybakov, S., Theis, F. J., Angerer, P., & Wolf, F. A. (2021). anndata: Annotated data. BioRxiv.

Wang, J., & Tontonoz, P. (2017). Pioneering EBF2 remodels the brown fat chromatin landscape. Genes and Development, 31(7), 632–633. 10.1101/gad.299644.117

Wang, W., Kissig, M., Rajakumari, S., Huang, L., Lim, H. W., Won, K. J., & Seale, P. (2014). Ebf2 is a selective marker of brown and beige adipogenic precursor cells. Proceedings of the National Academy of Sciences of the United States of America, 111(40), 14466–14471. 10.1073/pnas.1412685111

Wang, W., & Seale, P. (2016). Control of brown and beige fat development. Nature Reviews Molecular Cell Biology, 17(11), 691–702. 10.1038/nrm.2016.96

Wang, Y., & Sul, H. S. (2009). Pref-1 Regulates Mesenchymal Cell Commitment and Differentiation through Sox9. Cell Metabolism, 9(3). 10.1016/j.cmet.2009.01.013

Weiler, P., Lange, M., Klein, M., Pe’er, D., & Theis, F. (2024). CellRank 2: unified fate mapping in multiview single-cell data. Nature Methods, 21(7), 1196–1205. 10.1038/s41592-024-02303-9

Wolf, F. A., Angerer, P., & Theis, F. J. (2018). SCANPY: Large-scale single-cell gene expression data analysis. Genome Biology, 19(1). 10.1186/s13059-017-1382-0

Wolock, S. L., Lopez, R., & Klein, A. M. (2019). Scrublet: Computational Identification of Cell Doublets in Single-Cell Transcriptomic Data. Cell Systems, 8(4). 10.1016/j.cels.2018.11.005

Xu, Z., Wang, W., Jiang, K., Yu, Z., Huang, H., Wang, F., Zhou, B., & Chen, T. (2015). Embryonic attenuated Wnt/β-catenin signaling defines niche location and long-term stem cell fate in hair follicle. ELife, 4. 10.7554/elife.10567

Yong, L. W., Lu, T. M., Tung, C. H., Chiou, R. J., Li, K. L., & Yu, J. K. (2021). Somite Compartments in Amphioxus and Its Implications on the Evolution of the Vertebrate Skeletal Tissues. Frontiers in Cell and Developmental Biology, 9. 10.3389/fcell.2021.607057

Zheng, G. X. Y., Terry, J. M., Belgrader, P., Ryvkin, P., Bent, Z. W., Wilson, R., Ziraldo, S. B., Wheeler, T. D., McDermott, G. P., Zhu, J., Gregory, M. T., Shuga, J., Montesclaros, L., Underwood, J. G., Masquelier, D. A., Nishimura, S. Y., Schnall-Levin, M., Wyatt, P. W., Hindson, C. M., … Bielas, J. H. (2017). Massively parallel digital transcriptional profiling of single cells. Nature Communications, 8. 10.1038/ncomms14049

