## Supplementary material for "SOX9 is part of a combinatorial marker that reveals early development and embryological origins of the mouse brown adipose tissue depots": Table S1

**Supplementary Table 1:** Primary antibodies used in this study

| **Antigen** | **Antibody source and cat#** | **Dilution** | **species** |
|---|---|---|---|
| EBF2 | R&D Systems AF7006 | 1:13 | sheep |
| GATA6 | Cell Signaling Technology 5851 | 1:200 | rabbit |
| Muscle Actin | Abcam ab156302 | 1:200 | rabbit |
| Muscle Myosin | Millipore Sigma M4276 | 1:250 | mouse |
| PDGFR$\alpha$ | R&D Systems af1062 | 15 µg/mL | goat |
| PECAM | Developmental Studies Hybridoma Bank 2H8 | 1:15 | Armenian hamster |
| PPAR$\gamma$ | Thermo-Fisher MA5-14889 | 1:200 | rabbit |
| RFP | Chromotek 5f8 | 1:1000 | rat |
| SOX9 | Millipore Sigma AB5535 | 1:500 | rabbit |
| SOX9 | R&D Systems AF3075-SP | 1:200 | goat |
| Tenascin | Millipore Sigma T3413 | 1:100 | rat |
